# Single-cell long-read sequencing of the experience-induced transcriptome

**DOI:** 10.1101/2025.09.12.674885

**Authors:** Sheridan Cavalier, Paul W. Hook, Richard L. Huganir, Winston Timp

## Abstract

Neural activity drives transcriptional events that are critical for learning. Activity-induced transcript isoform expression and alternative splicing are cell-type specific events typically obscured by sequencing approaches that restrict read length. We combined single-cell transcriptomics with Nanopore long-read sequencing to resolve these phenomena, generating the first dataset profiling learning-induced gene and isoform expression in individual mouse hippocampal cells.

Due to their length, the majority of reads we generated could be uniquely mapped to their respective gene *and* isoform features in the mouse reference. ∼20k cells from 16 mouse samples were sequenced and clustered on the basis of gene expression, yielding 21 hippocampal cell types. Differential expression analysis revealed 1,266 significantly experience-variable isoforms, revealing novel splicing behavior in many synaptic genes.

This work is the first to comprehensively profile activity-induced isoform expression, demonstrating that single cell long-read sequencing can reveal new layers of transcriptional complexity.

## INTRODUCTION

Cells respond to environmental signals with differential gene expression and alternative splicing^1,2^, producing signal-specific RNA transcripts that influence cell behavior and biology. This is perhaps most remarkable in the brain, where neurons respond to activity by producing an array of functionally specialized and often cell-type specific transcripts^3–6^. The resulting activity-induced transcriptome and its downstream effectors modulate synaptic potentiation^7^, neural plasticity^8,9^, memory consolidation^10^ and behavior^11^.

Foundational studies of activity-induced gene expression in brain tissue report the involvement of hundreds of genes^12–14^ ranging in function from synaptic transmission^15^ to epigenetic regulation^16^. Not only does this transcriptional response recruit a diverse ensemble of genes, but these genes also differ largely across brain cell type^17–19^ and activity state^20,21^. Genes expressed in the brain are also subject to activity-induced alternative splicing, resulting in differential isoform expression^22^. For example *Bdnf*, which facilitates synaptic potentiation, responds to neural activity with alternative promoter usage^23,24^. Similarly, the activity-induced alternative poly-adenylation of *Homer1* generates a form of the encoded scaffold protein (Homer1a) that modulates synaptic scaling^3,25^.

Despite the demonstrated diversity of the activity-induced transcriptome across cell types and transcript isoforms, characterizing this response at both single-cell and isoform resolution has been technologically difficult. 10X Genomics Chromium, a droplet-based single-cell RNA-seq (scRNA-seq) platform, was developed for short-read sequencing^26^, limiting analysis to gene-level expression. In contrast, long-read Nanopore (ONT) sequencing generates reads from single, full length molecules that span multiple exon junctions and can be assigned to isoform features in the reference transcriptome^27–29^. Inspired by groups that have successfully combined single-cell transcriptomics with long-read sequencing^30,31^, we generated single-cell isoform expression data from mouse hippocampal samples using 10X Genomics and ONT. The resulting isoforms from each cell were quantified and analyzed for differential expression.

To specifically profile the activity-induced transcriptome with single-cell isoform sequencing, we applied this method to contextual fear conditioning, a behavioral learning paradigm that induces differential gene expression in the hippocampus^32–35^. We performed differential expression analysis on hippocampal dissociations from fear-conditioned, unconditioned, and naive mice to identify genes and isoforms that are modulated by these contextual experiences. Roughly 20% of the experience-variable isoforms we detected arose from alternatively-spliced genes that showed no differential expression at the gene level, suggesting these differences would be invisible without isoform resolution. With these expression data we were able to resolve experience-induced transcriptome and isoform feature-level differences within single hippocampal cells.

## RESULTS

### Single-cell long-read sequencing with 10X Genomics and ONT generates gene expression data comparable to short-read

To improve accuracy and enable demultiplexing of scRNA-seq reads from ONT R9.4 flowcells, we used a modified version of the Rolling Circle Amplification to Concatemeric Consensus (R2C2) method^36^. This method uses rolling circle amplification on circularized cDNA to generate long, repeat-concatemer molecules for sequencing and linear consensus calling. Using R2C2 on 10X-barcoded cDNA from mouse hippocampal dissociations (n=16), we were able to demultiplex ∼**94%** of barcode-containing molecules sequenced on ONT’s PromethION (**Table S1**).

Encouraged by our ability to demultiplex cell barcodes from ONT reads, we then wanted to ensure that we generated enough reads per cell to yield interpretable gene expression information. To validate that long-read scRNA-seq can yield the same gene expression information as canonical short-read scRNA-seq, we generated a 10X cDNA library targeting **2**,**000 cells** from one mouse hippocampal sample and divided it for sequencing with Illumina and R2C2+ONT (**Fig 1A**).

**Figure 1:**
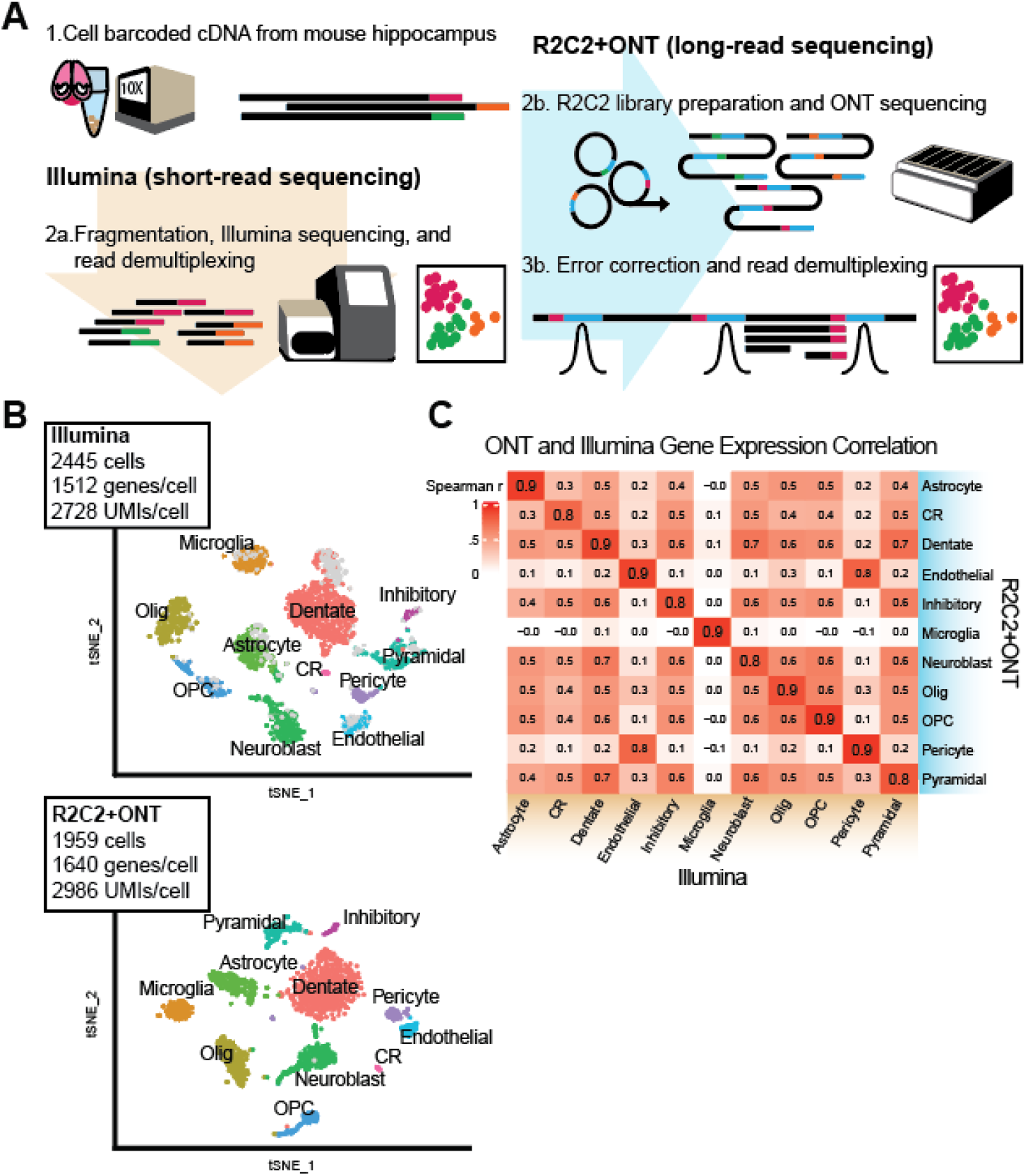
**A**. Schematic of Illumina (orange) and R2C2+ONT (blue) sequencing workflows used to generate two different scRNA-seq libraries from the same mouse hippocampal dissociation. **B**. tSNE plots generated from Illumina (top) or ONT (bottom) sequencing of the same hippocampal single-cell dissociation. Clusters are labeled by assigned cell type. Gray points on the Illumina plot denote platform-exclusive barcodes (i.e. found only in Illumina). **C**. Pairwise Spearman correlation values between gene expression profiles from Illumina (columns) and ONT (rows) cell types. Correlation tests were performed using the average cell-type expression values of the top 1000 most variable hippocampal genes.

Reads were mapped to the mouse GRCm39 reference genome, demultiplexed, and collapsed using Cell Ranger for Illumina (**Table S1**) or our custom analysis pipeline (https://github.com/SheridanCavalier/2025_sc_iso_paper) for the ONT data. To determine the unique molecular identifier (UMI) threshold at which a barcode is considered a cell (above background) in the ONT data, we imposed the same percentile-based criteria Cell Ranger uses for Illumina (Methods). For each library (Illumina or ONT), cells were clustered by gene expression with Seurat V3^37^. The resulting cell-type clusters in each dataset were assigned using canonical expression markers^38,39^ (**Fig 1B** and **Table S2**). We found that **99.59%** (1951) of cells recovered in the ONT dataset were also present in the Illumina dataset. The cells recovered exclusively by Illumina (494 cells) were present as barcodes in the long-read dataset; however, they fell short of the UMI threshold needed to qualify them as cells (**Fig S1**).

To confirm that cell-type gene expression is comparable between the two platforms, we examined the consistency of cell-type assignment between the two datasets. Across the 11 cell types, **96.45%** of ONT barcodes had the same cell-type assignment in the Illumina data (**Table S3**). When comparing each cell type’s expression of the top 1000 most variable hippocampal genes, every ONT cell type was most highly correlated with the matching Illumina type (**Fig 1C**).

### ONT sequencing of 10X samples generates single-cell isoform expression data

Reassured by gene-level agreement between Illumina and ONT, we examined the transcript isoform resolution of each dataset. The average insert length of cell-derived ONT reads in our dataset was **882 nucleotides** (nt), nearly ten times that of our short reads which were the standard length for 10X Chromium (92 nt) (**Fig 2A**). We reasoned that longer reads would allow for more direct isoform identification than the short-read standard, which in the 10X 3’ Gene Expression protocol is confined to the final 92 nucleic acids of each mRNA.

**Figure 2:**
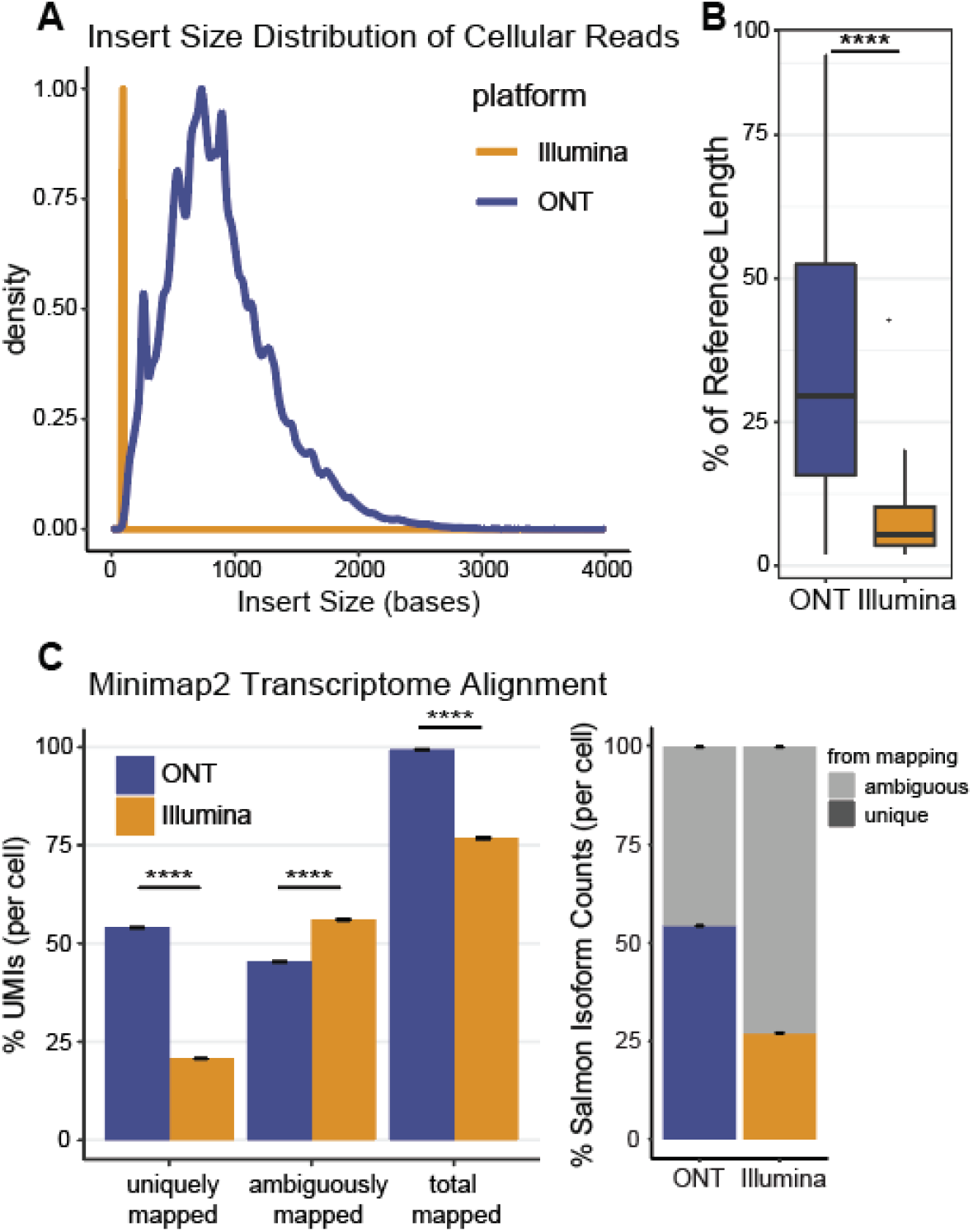
**A**. Insert size distributions of Illumina (orange) or ONT (blue) cDNA reads from the same hippocampal 10X scRNA-seq library. **B**. Percentage of reference transcript length covered by transcriptome-aligned ONT (blue) or Illumina (orange) reads. ONT reads covered a significantly greater percentage of their assigned reference transcript (median=38.9%) than Illumina reads (median=4.6%) as determined by a Mann-Whitney U Test (p-value < 2.2e-16). **C**. Left panel: Results from Minimap2 alignment of reads to the reference transcriptome (GENCODE M32). 54.0% of ONT reads mapped uniquely to one reference isoform compared to 20.7% of Illumina reads (Welch’s t, p< 2.2e-16). A further 45.3% of ONT and 56.0% of Illumina reads mapped to multiple reference isoforms (Welch’s t, p< 2.2e-16). The average mapping rate per cell was 99.3% in ONT and 76.7% in Illumina (Welch’s t, p< 2.2e-16). Right panel: Uniquely and ambiguously-mapped reads normalized by total mapped to reflect the proportion of isoform counts from Salmon that were observed (unique) vs inferred (ambiguous).

We evaluated isoform resolution in each dataset by first aligning cellular reads from Illumina and ONT to the mouse reference *transcriptome* (GENCODE, release M32) using Minimap2^40^. For each primary alignment, we compared the observed read length to the length of the reference isoform it aligned to (**Fig S2**). We found that the median ONT read was 38.9% of its reference isoform length, significantly greater than the 4.6% calculated for our transcriptome-aligned Illumina reads (Mann-Whitney, p-value < 2.2e-16) (**Fig 2B**). Transcriptome alignment files from each cell were then quantified with Salmon^41^. Salmon derives isoform abundance estimates from uniquely mapped reads to assign and quantify ambiguously mapped reads (i.e. those aligning to multiple reference isoforms). Of the average 99.4% of ONT reads per cell that mapped to the transcriptome, 54.0% uniquely aligned to a single isoform in the reference (**Fig 2C**, left panel). This was over twice the amount of uniquely mapped reads as the Illumina data (20.7%), in which isoform counts had to be largely inferred from ambiguously mapped reads (**Fig 2C**, right panel).

### Long-read scRNA sequencing reveals experience-variable gene expression

Activity-induced gene expression occurs in multiple cell types and across induction stimuli. In the hippocampus, a brain region involved in encoding contextual memory, activity induces waves of differential gene expression, including immediate early genes (IEGs) *Fos, Egr1*, and *Junb*^32,42-47^. *Fos* is induced in only a subset of responsive cells and is a marker of recently-activated “engram” neurons^14,48-51^. The cell autonomous expression of *Fos* emphasizes the experimental need for single-cell resolution, even within the same cell type. Single-cell RNA sequencing has further demonstrated the extreme cell-type heterogeneity of the activity-induced transcriptome within and across brain regions^18,19^. To characterize activity-induced gene and isoform expression in hippocampal cells, we applied single-cell long-read sequencing to contextual fear conditioning.

Contextual fear conditioning integrates novel exposure and classical learning–allowing us to study transcriptional responses to three different contextual experiences: Conditioned, Unconditioned, and Naive. Fear-conditioned (C+) animals (n=6) were exposed to a novel context for five minutes while simultaneously receiving three 0.5 mA foot shocks, Unconditioned (U+) animals (n=6) were exposed to the novel context for five minutes in absence of foot shocks, Naive (N) animals (n=4) were not exposed to the context. Half the context-exposed mice (C+ and U+) were sacrificed at 10 minutes and half at 60 minutes post-exposure to capture the early experience-induced transcriptional response (**Fig 3A** and **Table S4**).

**Figure 3:**
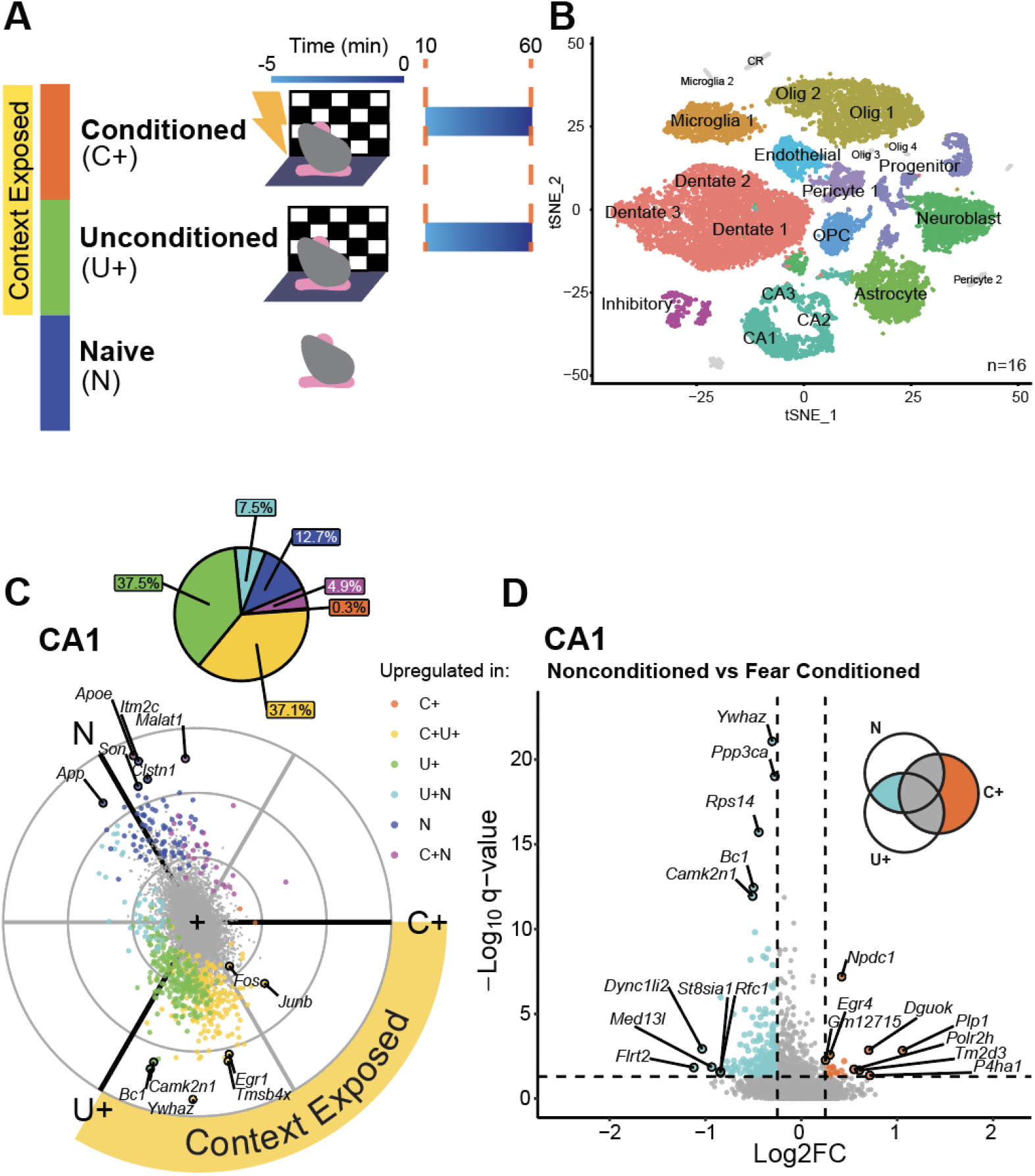
**A**. The three contextual experiences: Conditioned (C+) mice (orange) were exposed to the novel context for 5 min during which three foot-shock stimuli were prescribed and then sacrificed at 10 or 60 min following context-exposure. Unconditioned (U+) mice (green) were exposed to the novel context for 5 min and sacrificed 10 or 60 min following. Naïve (N) mice (blue) received no context exposure. **B**. tSNE of dorsal hippocampal cells from 16 mice across three contextual experiences, clustered by gene expression. Cell type clusters in gray contain <100 cells and were excluded from downstream analysis. **C**. Radial expression plot of CA1 genes. A gene’s distance r and angle Ө from the origin represent its overall magnitude and direction of expression across the three contextual experiences. Significantly experience-variable genes (q value <.05, FDR ≤.05) are colored by the experience(s) in which they were upregulated (log2FC >.25). Top ten genes by r are labeled in addition to Fos and Junb. (inset) Percentage of total significant CA1 genes upregulated by each experience. **D**. Volcano plot of differentially-expressed genes between fear-conditioned and nonconditioned CA1 neurons. For each gene, expression in C+ cells was compared with the averaged expression in U+ and N cells, resulting in the log2 fold change values shown. Q values are from DESeq2 analysis. Top five genes per condition by significance and log2 fold change are labeled.

Whole-cell suspensions from each mouse were generated from acute dorsal hippocampal slices (Methods). Tetrodotoxin (TTX) and Actinomycin-D were included to minimize spontaneous activity. Single-cell suspensions were processed into barcoded cDNA using the 10X Genomics 3’ Gene Expression Kit v3.1. Unfragmented barcoded cDNA was circularized, amplified and prepared for ONT sequencing (Methods). Reads were processed, demultiplexed and collapsed using our custom pipeline (https://github.com/SheridanCavalier/2025_sc_iso_paper)(**Table** S1). Gene count matrices were generated by mapping reads to the mouse GRCm39 reference genome and quantifying primary alignments with FeatureCounts^52^. Cells were then filtered, clustered by their gene expression profiles, and assigned (**Table S5**). The resulting single-cell RNAseq dataset of the dorsal hippocampus encompasses 21 cell types, three different contextual experiences, and two time-points (10 or 60 minutes post-exposure). We chose to combine time points to focus our analysis on contextual experience rather than the more subtle axis of time. Cell-type clusters with less than one-hundred cells (CR, Microglia 2, Olig 3, Olig 4, and Pericyte 2) were excluded from downstream analysis (**Fig 3B**). We observe no confounding of cell type clustering with exposure, mouse sex, or batch (**Fig S3**). We analyzed each cell type separately for experience-variable gene expression, using DESeq2’s likelihood ratio test (LRT) as recommended^53,54^. This analysis evaluates the relationship between contextual experience and each gene’s expression variance using linear modelling. A gene was considered significantly experience-variable (p <.05, FDR ≤.05) if its variance was better explained by including contextual experience as a term in the linear model. To control for sex and batch, we incorporated these terms as fixed effects in our model equations (Methods). This analysis yielded a combined 5,870 experience-variable genes (3,061 unique genes) across 14 cell types. To classify variable genes, we evaluated the log2 fold change of each pairwise comparison between the three experiences (i.e. C+ vs U+, C+ vs N, U+ vs N). A gene was considered upregulated by an experience if the log2 fold change for one or both pairwise comparisons was >.25 (**Supp Data 1**).

Oligodendrocytes, dentate neurons and CA1 pyramidal neurons respond to contextual experience with the greatest number of genes (a total 4,348) (**Fig S4**). Gene ontology (GO) enrichment analysis of these genes^55–57^ yielded expected shared (i.e. “Neuron Projection Development”) and distinct (i.e. “Myelination” in oligodendrocytes) functional responses to contextual experience from these three cell types (**Fig S5**).

To visualize overall gene expression across the three experiences for a given cell type, we generated radial plots inspired by volcano3D^58^. From our count matrices we calculated the average expression of each gene in each experience. We assigned each experience a fixed angle *θ* (C+=0°,U+=240°,N=120°) so that any gene’s expression could be described by three polar vectors (*r, θ*), where *r* is the average expression in an experience and *θ* is the angle assigned to that experience. The sum of these vectors generates the final placement of the gene on the radial expression plot, where *r* and *θ* represent overall magnitude and direction of expression (**Fig 3C**).

We first examined experience-induced gene expression in CA1 pyramidal cells, excitatory neurons that are activated by contextual experience and are critical for contextual memory consolidation and recall^47,59^. *Fos, Egr1* and *Junb* are activity-induced IEGs previously shown to be upregulated in CA1 neurons following novel context exposure or contextual fear conditioning^45,47,60,61^. We found *Fos, Junb*, and *Egr1* significantly upregulated in CA1 neurons by the unconditioned (U+) and fear-conditioned (C+) experiences, both of which involve novel context exposure (**Fig 3C** and **Fig S6**). To identify genes that most distinguish contextual learning (C+) from the other two experiences in the CA1, we created volcano plots from the log2 fold difference between C+ and averaged U+ and N gene expression values. We found *Plp1*, a gene encoding a critical component of myelin (Plp1)^62^ and also expressed in hippocampal neurons^63^, was most upregulated in C+ relative to U+/N CA1 cells (**Fig 3D**).

As mentioned, *Fos* is induced in a subset of activated neurons, the “engram”, that is responsible for encoding and maintaining a memory of the inducing stimulus^14,48-51^. We reasoned we could identify putative CA1 engram neurons in our dataset on the basis of *Fos* expression. Based on previous studies reporting ∼ 25% of the CA1 is activated by context exposure^64^, we set a normalized *Fos* expression threshold categorizing 21% of fear-conditioned (C+) CA1 neurons as engram cells (**Fig 4A** and **Fig 4B**). We then compared their gene expression to non-engram CA1 cells and found that genes most upregulated in the engram population are shared between C+ and U+ (**Figs 4C** and **4D**), suggesting that the CA1 engram encodes specifically the contextual element of the experience, which is common to both the fear-conditioned and unconditioned paradigms.

**Figure 4:**
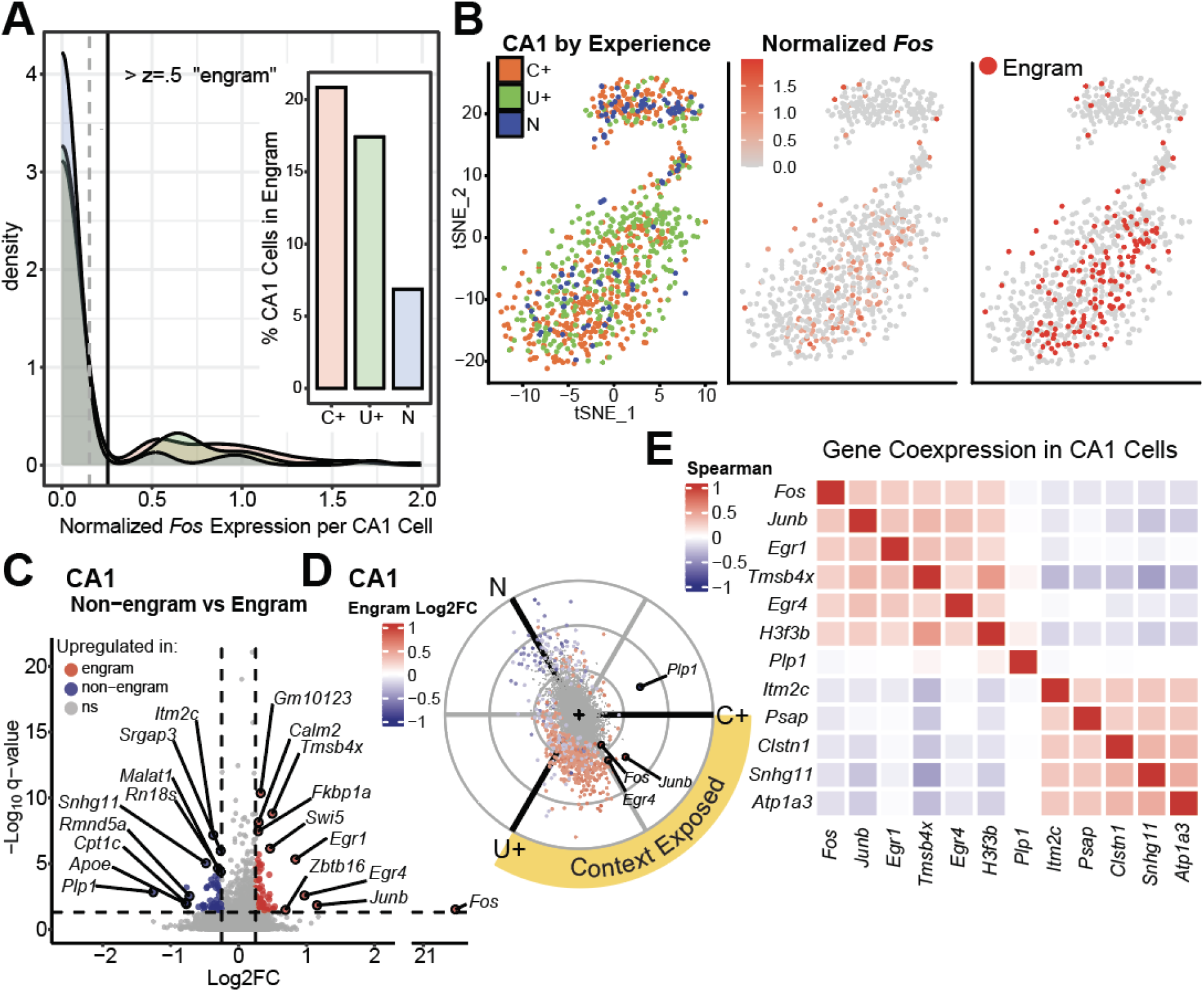
**A**. Distributions of normalized Fos expression per CA1 cell colored by experience. Mean and z-scores are based on the cumulative count distribution across all experiences. Cells with normalized Fos greater than.5 standard deviations (z >.5) above the mean were considered engram cells. (inset) Percentage of CA1 cells in the engram, by experience. **B**. CA1 cells clustered by experience-variable genes: colored by experience (left), depth-normalized Fos expression (middle), and engram cell identity (right). **C**. Volcano plot of pairwise comparison between CA1 engram and non-engram gene expression. **D**. CA1 gene radial plot colored by log2 fold change in C. Genes with |log2FC| > 1 are labeled. **E**. Gene co-expression in CA1 cells. Spearman correlation matrix between engram-variable genes, sorted by correlation r with Fos. Top five pos and neg genes are shown alongside Fos and Plp1.

Surprisingly *Plp1*, although strongly upregulated by fear conditioning (**Fig 3D**), was downregulated in engram cells**(Fig 4C**). Examining the co-expression of engram-variable genes across all individual CA1 neurons revealed that *Fos* and *Plp1* belong to different gene expression programs (**Fig 4E**). The upregulation of *Plp1* in fear-conditioned but not engram cells alludes to a population of CA1 neurons that is strongly affected by fear conditioning but is responding separately from engram neurons. Of the 250 genes upregulated by fear conditioning in CA1 neurons, 87.6% (219) were also upregulated in the unconditioned experience (**Fig 3C**). These results highlight transcriptional similarity between conditioned and unconditioned CA1 neurons during the early (< 1hr) exposure-induced response. This was not simply a failure to condition: in a separate experiment, fear-conditioned (C+) mice spent more time freezing compared to unconditioned (U+) littermates when re-introduced to the context four days later (**Fig S7**). We suggest the large overlap between C+ and U+ CA1 transcriptomes is indicative of the early activity-induced response to novel context exposure.

Compared to CA1 neurons, oligodendrocytes had a greater proportion of genes upregulated exclusively by fear conditioning (4.4%) (**Fig S8**). Oligodendrocytes are myelinating glial cells responsive to neural activity^65,66^ and experience^18^. We found that fear conditioning upregulated IEGs *Nfkbia*^67^ and *Sgk1*^68^ (**Fig S9**) in this cell type; the protein products of both genes are involved in NF-kappaB signalling^69^, which is implicated in hippocampal plasticity^70^. *Sgk1* has previously been identified as responsive to visual stimulation in mouse cortical oligodendrocytes^18^, suggesting alongside our results a common experience-induced gene expression program in oligodendrocytes across brain regions and stimuli. These differential gene expression data provide insight into cell-type specific responses to activity and allow for the first large-scale interrogation of gene expression across three distinct contextual experiences in the hippocampus.

### Long-read scRNA sequencing reveals cell-type and experience-variable isoform expression

Genes in the brain undergo activity-induced alternative splicing, resulting in differential isoform expression^22^. Although extremely valuable, *gene* count data are largely insufficient to detect *isoform* expression changes. While isoforms arising from activity-induced alternative splicing have been identified^3,6,23^, these findings are often derived from qPCR^6^ or FISH labeling^3^ of targeted isoforms from genes of interest, preventing broad profiling of isoforms without *a priori* knowledge. Alternatively, scRNA-seq is an ideal tool for whole transcriptome profiling and discovery; however, short-read scRNA-seq is limited to gene expression, precluding measurement of transcript isoforms. We leveraged our unique, single-cell long-read dataset to broadly and agnostically profile activity-induced isoform expression across hippocampal cell types.

To evaluate differential isoform expression in the ONT data, we first analyzed naive mice (n=4) for isoforms that were significantly variable between hippocampal cell types. Again we used the LRT built into DESeq2, this time to evaluate the influence of cell type on each isoform’s expression variance. We detected 8,908 unique transcript isoforms (16.1% of our hippocampal transcriptome) with significantly variable expression between at least two of the 14 cell types analyzed (**Supp Data 2**). For example, two of the cell-type variable isoforms detected, *S100a16-202* and *S100a16-201*, belong to *S100a16*, which encodes a calcium binding protein involved in calcium homeostasis and cognition^71^. We found astrocytes have significantly higher expression of *S100a16-202* (Wilcoxon Rank Sum = 98690, p-value = 2.739e-06) than oligodendrocytes and significantly lower expression of *S100a16-201* (Wilcoxon Rank Sum = 58515, p-value < 2.2e-16) (**Fig 5A**). These isoforms have different transcription start sites (TSS), resulting in alternative 5’ untranslated region (UTR) usage between them. This difference is detectable in the long-read data, but is unresolvable with short reads (**Fig 5B**). These data exemplify the ability of single-cell long-read sequencing to reveal differential isoform expression, and with pre-existing computational tools.

**Figure 5:**
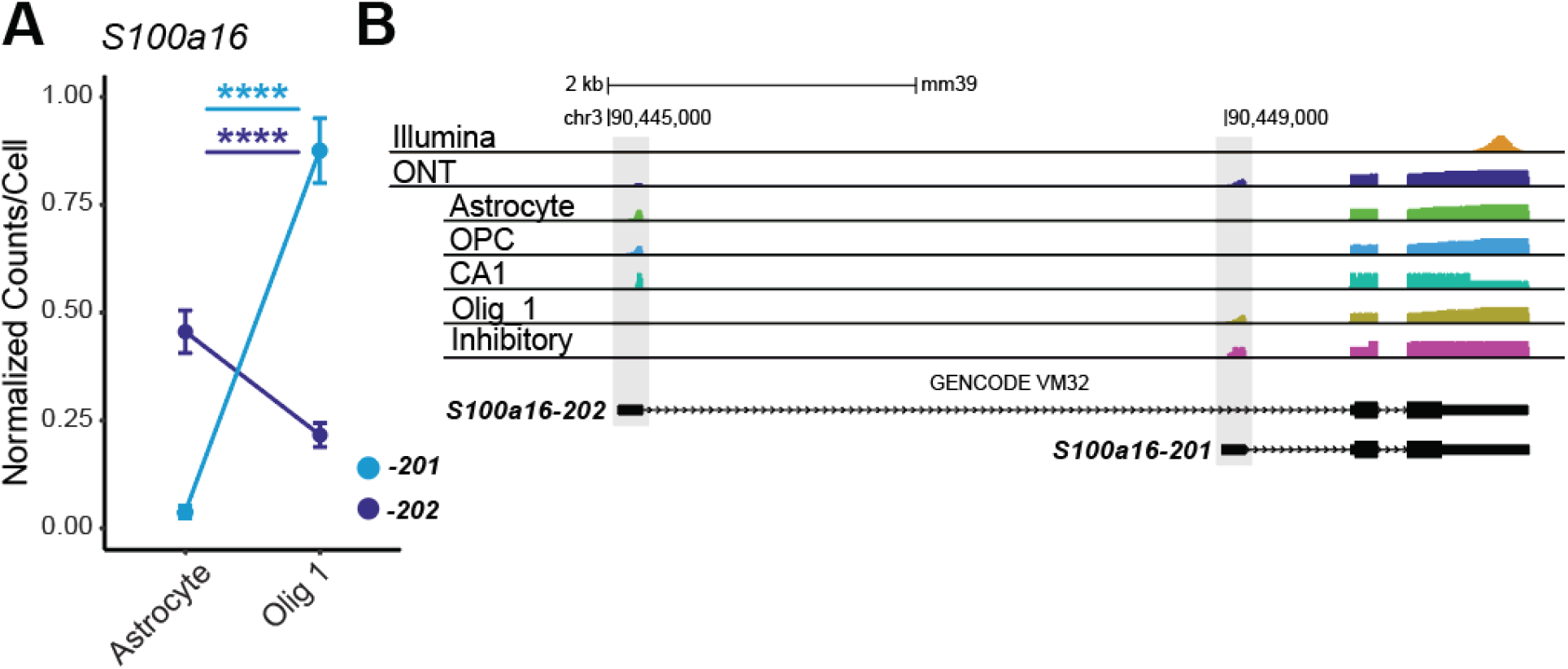
**A**. Depth-normalized counts per cell of isoforms S100a16-202 and S100a16-201 in astrocytes and oligodendrocytes. Astrocytes have significantly higher expression of S100a16-202 (W = 98690, p-value = 2.7e-6) than oligodendrocytes and significantly lower expression of S100a16-201 (W = 58515, p-value < 2.2e-16) as determined by a Wilcoxon Rank Sum test. **B**. Coverage histogram tracks for Illumina (orange) or ONT (blue) reads aligning to S100a16, visualized with UCSC Genome Browser. ONT reads are split into cell type tracks. Each track is scaled to its own coverage min and max for easier cross-track comparisons of alternative exon usage. Grey areas highlight the first exons (containing the 5’UTRs) of the isoforms shown.

To profile experience-variable isoform expression we applied the same differential expression analysis framework as for experience-variable genes, but now using the *isoform* count matrices obtained by quantifying transcripts with Minimap2 and Salmon (Methods). We separately evaluated each cell type in our dataset for experience-variable transcript isoforms using the LRT in DESeq2. This yielded a combined 1,972 experience-variable transcript isoforms (1,266 unique isoforms) across 14 cell types and belonging to 1,217 genes (**Supp Data 3**). Again we performed pairwise comparisons between experiences to further classify isoforms by the experience(s) that upregulated them (q<.05, FDR ≤.05, Log2FC >.25). With the resolution in our data we were able to identify the exact *Fos* isoform that is upregulated by context exposure, the only isoform with a defined coding sequence, *Fos-201* (**Fig S10**).

To better understand experience-induced alternative splicing, we classified experience-variable isoforms by the splicing behavior of their genes (**Fig 6A**). Single-isoform genes (giving rise to 15% of experience-variable isoforms) only express one isoform in a cell type. The majority (53.6%) of experience-variable isoforms arise from alternatively spliced genes that are themselves differentially expressed in the gene data. Perhaps most interesting are the 21.3% of experience-variable isoforms across cell types that arise from genes with *no* significant differential expression (**Fig 6A**). In these cases, differential expression of isoforms across experiences is cancelled out by gene-level count aggregation. These experience-induced alternative splicing phenomena, which produce a fifth of the activity-induced hippocampal transcripts in our data, are uniquely resolved with isoform resolution.

**Figure 6:**
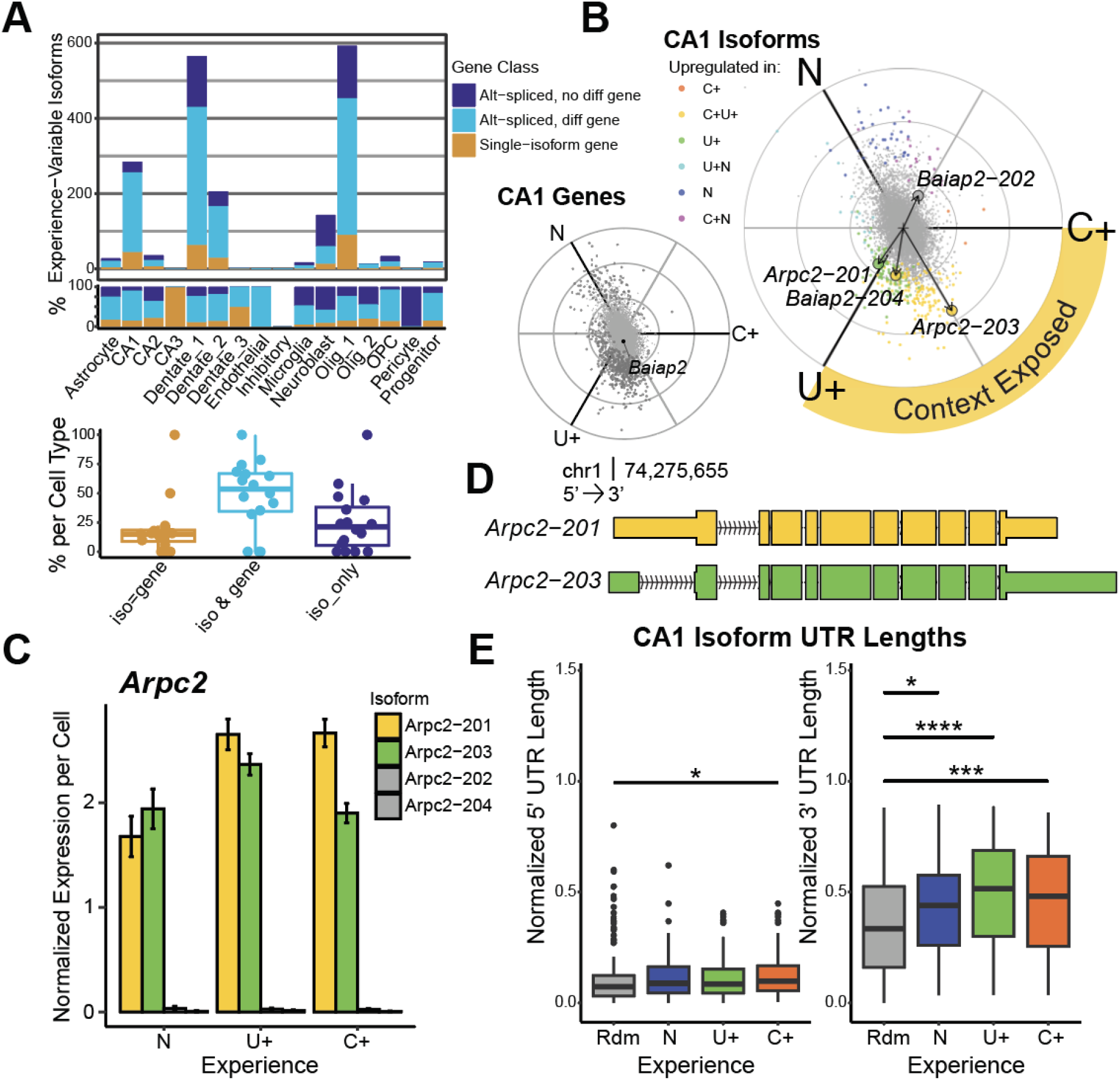
**A**. (top) Experience-variable isoforms per cell type, colored by gene class: Single-isoform genes (orange) express one isoform. Isoforms from alternatively-spliced genes are classified by whether the associated gene is (light blue) or is not (dark blue) also differentially expressed. (bottom) Boxplot of percent gene class per cell. A median 21.3% of experience-variable isoforms arise from genes that are not differentially expressed (iso_only). **B**. (left) Radial expression plot of CA1 genes. Gene Baiap2 is labelled. (right) Radial expression plot of CA1 isoforms. Isoforms Baiap2-204, Baiap2-202, Arpc2-201, Arpc2-203 are labelled. **C**. Depth-normalized expression of Arpc2 isoforms in CA1 cells, with Arpc2-201 (yellow) and Arpc2-203 (green) highlighted. **D**. Arpc2-201 (yellow) and Arpc2-203 (green) transcript models (GENCODE vM32). **E**. UTR lengths of experience-variable isoforms (N, U+, C+) and a random selection of 150 CA1-expressed isoforms (Rdm), normalized to transcript length. (left) Normalized 5’UTR lengths (Kruskal-Wallis, χ^2^=9.52, df=3, p=.023). (right) Normalized 3’UTR lengths. (Kruskal-Wallis, χ^2^=32.84, df=3, p=3.48e-7). Significance bars reflect pairwise Wilcoxon test results.

One example is *Baiap2*, which encodes one of the most abundant scaffolding proteins (Irsp53) in the postsynaptic density^72,73^. We discovered experience-induced alternative splicing of *Baiap2* in CA1 pyramidal neurons. Due to outcompeting expression of isoforms *Baiap2-204* and *Baiap2-202*, we could not detect differential expression of gene *Baiap2*, exemplifying the need for transcript resolution (**Fig 6B**). Context exposure induces *Baiap2-204* in CA1 neurons (**Fig 6B**). This transcript encodes a protein isoform of Irsp53 that lacks the canonical C-terminal PDZ binding motif (VSTV) required for association with the postsynaptic density^74^ (**Fig S11**). Exclusion of the PDZ binding motif by alternative splicing has previously been characterized in *Syngap1*^75,76^, which dissociates from the postsynaptic density in response to neural activity to facilitate synapse-specific potentiation^77^. While we observed a modest induction of *Syngap1-204* (lacking the PDZ binding motif) in context-exposed CA1 cells, this finding was not significant (**Fig S12**). Still, the similarities in subcellular localization and function between Baiap2 and Syngap1 proteins suggest the activity-induced alternative splicing we observe in *Baiap2* might serve a similar purpose of priming the synapse for potentiation.

Differential isoform expression analysis also revealed two experience-regulated isoforms of *Arpc2. Arpc2* encodes the actin-binding subunit of Arp2/3 and facilitates the cytoskeletal reorganization required for dendritic development^78,79^. We observed that these isoforms are differentially regulated by contextual learning in CA1 neurons. While *Arpc2-201* and *203* are both upregulated in the unconditioned experience (U+), fear conditioning (C+) induces the expression of isoform *Arpc2-201* exclusively (**Fig 6 B & C**), which harbors the same coding sequence as *Arpc2-203* but has a comparatively longer 5’ and shorter 3’ UTR (**Fig 6D**).

After observing experience-regulated alternative UTR usage in *Arpc2*, we quantified UTR lengths for all experience-variable isoforms in the CA1. To provide greater context for interpretation, we compared the UTR lengths of significant isoforms to those of an equivalently-sized random selection of CA1-expressed transcripts (Rdm) (**Fig 6E**). Comparing the UTR lengths of protein-coding CA1 isoforms revealed significantly longer 5’ UTRs in learning-induced (C+) isoforms relative to the average (Rdm) (Wilcoxon Rank Sum=8888, p=.003).

More generally, the 3’UTRs of experience-variable isoforms were significantly longer than average (Rdm) regardless of experience (Kruskal-Wallis, χ^2^=32.84, df=3, p=3.48e-7), with 3’UTRs in unconditioned (U+) isoforms being the longest. Thus, experience-variable CA1 isoforms generally seem to reflect the behavior of *Arpc2*, in which learning-induced (C+) isoforms harbor comparatively longer 5’UTRs and unconditioned (U+) isoforms comparatively longer 3’ UTRs.

To evaluate engram-variable isoform expression, we analyzed our previously defined subset of *Fos*+ CA1 engram cells (**Fig 4A & B, Fig 7A**). We found that IEG isoforms *Egr1-201, Junb-201* and *Egr4-201* are upregulated in the engram population and *Plp1-201* is strongly downregulated (**Fig 7B**), similar to the gene data. We next calculated the coexpression of isoforms in individual CA1 cells. To focus on the isoform landscape strictly within engram cells, we chose to just highlight isoforms that were positively correlated with *Fos-201* (**Fig 7C**), including *Arpc2-201* which we discovered is strongly induced by context exposure. These data demonstrate the value of isoform resolution in individual cells.

**Figure 7:**
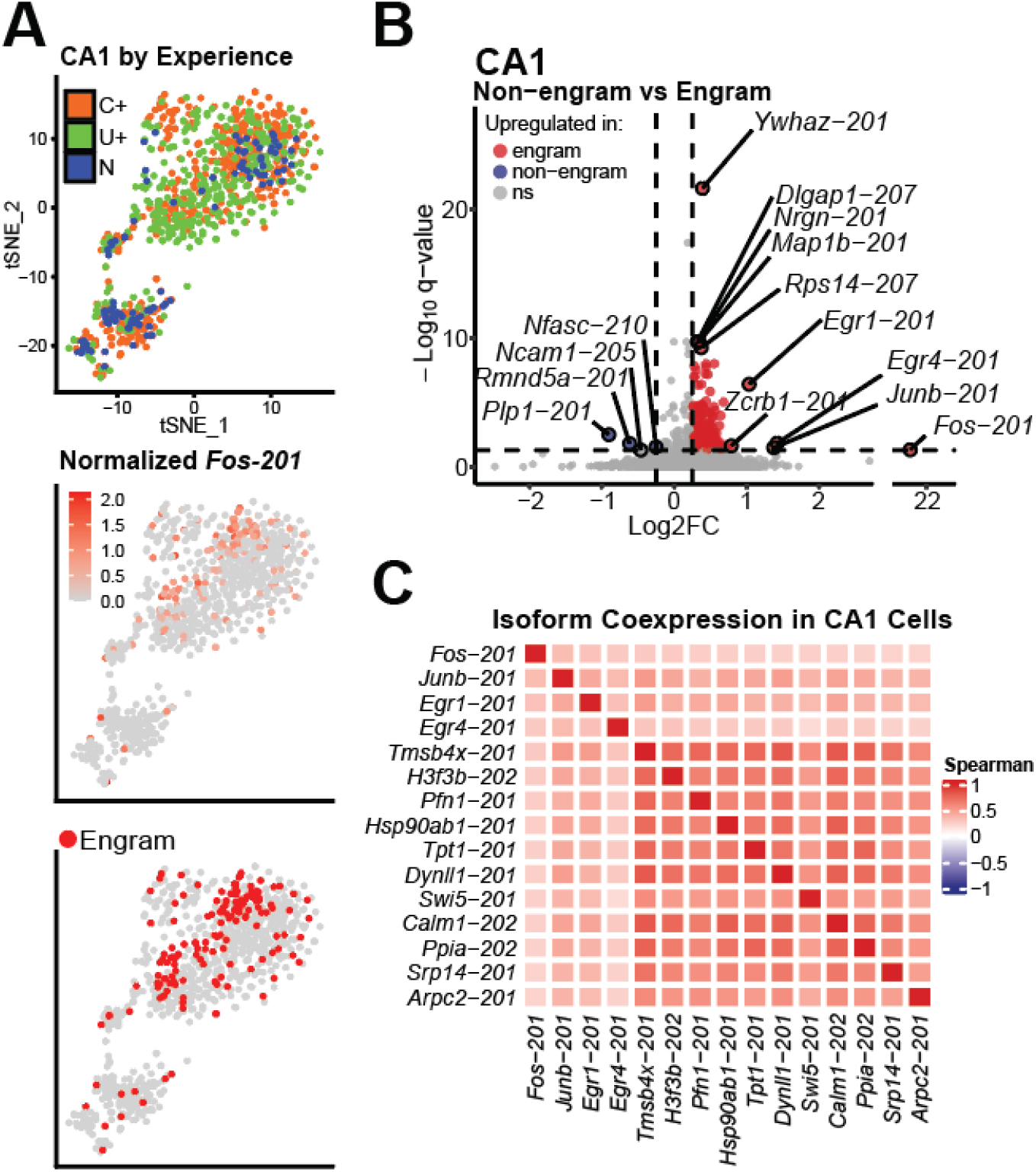
**A**. CA1 cells clustered by experience-variable isoforms: colored by experience (top), depth-normalized Fos-201 expression (middle), and engram cell identity (bottom). **B**. Volcano plot of pairwise comparison between CA1 engram and non-engram isoform expression. **C**. Isoform co-expression in CA1 cells. Spearman correlation matrix between engram-variable isoforms, sorted by correlation r with Fos-201, top 15 shown.

## DISCUSSION

Activity-induced transcription is a complex biological process that produces diverse cell type-specific gene and isoform responses^3,5,13,18,23^. While genes involved in this response have been well-studied, cell-type expression of isoforms has remained largely unexplored due to technical limitations. Short-read single-cell sequencing is largely limited to gene count data, while long-read bulk sequencing cannot capture cell-type heterogeneity. We combined 10X Chromium with ONT sequencing, to characterize cell-type gene and isoform responses to three different contextual experiences in the mouse hippocampus: fear-conditioned, unconditioned, and naive. We detected 3,061 unique experience-variable genes and 1,266 unique isoforms across 14 hippocampal cell types. Of these isoforms, over 20% have expression dynamics that are undetectable at the gene level, demonstrating the value of isoform resolution in characterizing cellular responses to learning and experience. This is the first dataset to our knowledge that characterizes learning-induced differential gene *and* isoform expression in single cells.

We first explored gene-level changes, finding context exposure-induced upregulation of *Fos* and other activity-responsive IEGs in CA1 pyramidal cells^13,24,42^. A significant advantage of single-cell analysis is the ability to detect differential expression even within cell type, which is not possible with sorting style methods such as flow cytometry. We chose to focus on *Fos+* CA1 cells because this engram population is most well-studied in the hippocampus^42,43,45,46,49^; however, engram behavior has been reported in multiple cell types and associated with a variety of IEG markers^80^. The genes most upregulated in *Fos+* CA1 engram neurons were largely shared between context-exposed animals, indicating that *this* engram population is encoding the context, which is common to both experiences. We also noticed a separate CA1 subpopulation of *Plp1*+ cells, anticorrelated with the engram population, but also strongly affected by fear conditioning. These data suggest that different subpopulations, even *within* cell type and experience, are uniquely responsive to learning, perhaps as a way of further delineating the engram subpopulation and encoding memory.

In oligodendrocytes we observed strong induction of *Sgk1* and *Nfkbia* specifically by fear conditioning (C+). These genes encode members of the NF-κB signalling pathway, which has been implicated in synaptic signalling^69^. *Sgk1* specifically has previously shown visual experience-induced expression in oligodendrocytes^18^, suggesting a more global engagement of this pathway across multiple brain regions and stimuli. That this response is elevated in oligodendrocytes alludes to a potential function of this pathway in regulating activity-induced myelination and neural connectivity.

When we look beyond gene expression, the hidden transcriptional differences are substantial; 20% of experience-variable isoforms arose from genes that had no differential expression. One example, *Baiap2*, a gene encoding the abundant synaptic protein Irsp53^73^, undergoes context exposure-induced alternative splicing in CA1 neurons to exclude the C-terminal PDZ-ligand sequence that is present in the naive protein. This novel splicing behavior, which is intractable without isoform resolution, is similar to the synaptic protein Syngap1^73,75,76^, alluding to a potentially shared mechanism of experience-induced alternative splicing. We also discovered learning-induced alternative transcription start site (TSS) usage in *Arpc2*, a gene that is important for synaptic structure and function^78,79^. Fear-conditioned (C+) CA1 neurons have a higher proportion of *Arpc2-201*, an isoform of the gene that harbors the same coding sequence but a longer 5’UTR than the isoform upregulated in unconditioned cells (U+), *Arpc2-203*. The expression of these isoforms confirms that, at the mRNA sequence level, there is a distinction between learning about a context and merely experiencing it and that this difference precipitates in some isoforms as alternative TSS usage. Although the activity-induced splicing properties of this gene were previously unknown, the long 5’UTR *ARPC2* isoform in humans has been shown to harbor an internal ribosome entry site^81^. When we evaluated the 5’ and 3’ UTR lengths of all experience-regulated isoforms in the CA1, we found that learning generally yields isoforms with longer 5’ UTRs, a result consistent with the *Arpc2* data. The learning-induced alternative TSS usage we observe is potentially a means of encoding stimulus activity within the regulatory sequence of resulting transcripts, likely influencing protein binding, trafficking and/or translation.

Further exploration of these isoforms, their putative sequence motifs, regulation and subcellular localization is needed to determine the specific effects of each of the thousands of experience-induced alternative splicing decisions revealed in our dataset. The greatest limitation of isoform quantification in single cells is the sparsity of the resulting count data, which we found greatly hampers the statistical power of differential isoform expression analysis. Deeper sequencing per cell or targeted enrichment for isoforms of interest are plausible ways to circumvent low count numbers; however, data sparsity is an unfortunate drawback of most single cell sequencing experiments. Fortunately, improvements in sequencing yield since we generated this data make single-cell long-read sequencing even more efficient and accessible. As long-read sequencing and single-cell transcriptomic methodologies continue to converge, we believe that isoform level resolution will quickly become the new standard for single-cell RNA sequencing.

## METHODS

### Mice

Experiments were conducted using C57BL/6J male and female mice (The Jackson Laboratory) ranging in age from 30 to 45 days old. Littermates were separated by sex and housed in pairs.

### Behavior

Each mouse was handled five minutes for five consecutive days prior to behavioral testing. Fear-conditioned (C+) mice (n=6) were placed in a sound-proofed footshock chamber with visual and olfactory contextual cues (checker-patterned wall and 70% ethanol) for five minutes. One 0.5 mA, 1s duration footshock was triggered at minutes 2, 3, and 4 of the conditioning paradigm for a total of three stimulations. Animals were sacrificed for single cell dissociation ∼10 min (n=3) or 1 hour (n=3) following fear conditioning. Unconditioned (U+) mice (n=6) were exposed to the described context for five minutes in absence of footshocks stimulation. Animals were sacrificed for single cell dissociation ∼10 min (n=3) or 1 hour (n=3) following context exposure. Naive (N) mice (n=5) were not exposed to the described context and were sacrificed for single cell dissociation ∼24 hours after their last handling.

Cage mates of C+ and U+ mice underwent the same conditioning and context exposure paradigms, but were returned to their home cages for recall testing four days later: mice were re-introduced to the same context for five minutes without footshocks and freezing time was manually scored to assess conditioned fear.

### Single cell dissociation

10 min or 1 hr following behavior, mice were euthanized with isoflurane and administered a transcardial perfusion of cold, carbogenated (95% O2, 5% CO2) low-activity ACSF (25 mM Glucose, 20 mM HEPES, 1.25 mM NaH2PO4, 25 mM NaHCO3, 2.5 mM KCl, 96 mM NMDG, 3 mM myo-inositol, 12 mM N-acetyl-cysteine, 5 mM ascorbic acid, 3 mM Na pyruvate,.01 mM taurine, 2 mM trehalose, 10 mM MgCl2,.5 mM CaCl2) at pH=7.4 and mOsm=290. We included 5ug/mL Actinomycin D and.1uM Tetrodotoxin in the perfusion buffer to minimize spontaneous activity and block further transcription. The brain was extracted and mounted on a vibratome stage (Leica VT1200) for acute coronal slicing of the dorsal hippocampus. Four, 300um thick coronal slices were taken from each hemisphere and the dorsal hippocampus was manually dissected from each slice. The dissected hippocampi were digested in room-temperature, carbogenated digestion buffer (low-activity ACSF with 5ug/mL Actinomycin D,.1uM Tetrodotoxin, and 1mg/mL pronase) for 70 minutes.

Following digestion, hippocampi were transferred to an Eppendorf tube and excess digestion buffer was removed and replaced with cold, carbogenated low-activity ACSF. Tissue was manually triturated with fire-polished Pasteur pipettes. Large debris was allowed to settle and the suspension was pipette-transferred to another Eppendorf tube and pelleted in a microcentrifuge for 5 min at 300 rcf. Supernatant was removed and the pellet was resuspended with a pipette in cold, carbogenated resuspension buffer (25 mM Glucose, 20 mM HEPES, 1.25 mM NaH2PO4, 30 mM NaHCO3, 2.5 mM KCl, 92 mM NaCl, 1 mM MgCl2, 1 mM CaCl2) at pH=7.4 and mOsm=290. The resulting solution was filtered through a 40um cell strainer. Cell count and health were assessed with a hemocytometer.

### Library preparation and sequencing

Single-cell suspensions were processed with the 10X Chromium 3’ Gene Expression kit (v3.1) into barcoded cDNA. We diluted each sample with resuspension buffer to target 2,000 cells for recovery. For Illumina sequencing the 10X protocol was followed as directed. For ONT sequencing, the barcoded and amplified cDNA from Step 2 of the protocol was eluted prior to fragmentation and prepared for sequencing with R2C2.

Unfragmented, barcoded cDNA was circularized and amplified by phi29 as described^36^. After amplification, branched cDNA was digested into linear concatemers and sheared to 30kb (Diagenode Megaruptor 2). After shearing, long cDNA was purified using the Short Read Eliminator XS kit (Pacific Biosciences). Shearing and purification allowed us to standardize library size and extend the life expectancy of ONT sequencing pores. These libraries were prepared for sequencing using the SQK-LSK110 kit from ONT. Illumina sequencing was performed on the MiSeq device (MiSeq Reagent Kit v3, 150 cycles). ONT sequencing was performed on the PromethION device (R9.4.1, FLO-PRO002).

### Read processing and cell type clustering

Fast5 files from PromethION sequencing were basecalled with Guppy (ONT, version 5.0.12). Resulting reads were processed with C3POa: the subreads in each concatemeric read were identified by partial order alignment, aligned to each other using Minimap2 (v2.17) and polished with Racon (v1.4.19)^82^ to generate a consensus read.

We used a custom analysis pipeline (https://github.com/SheridanCavalier/2025_sc_iso_paper) for demultiplexing and UMI-collapsing ONT scRNA-seq reads from each sample. Consensus reads were oriented and trimmed so the last 28 nt contain the 10X UMI and barcode. Using the 10X barcode whitelist, reads were demultiplexed first by exact matching. These barcodes made up an abridged list against which the remaining unmatched reads were demultiplexed with an error tolerance of 2 nt (L < 3). Demultiplexed reads were then aligned to the mouse reference genome (GRCm39, June 2020) using Minimap2 with the parameters: `--secondary=no -ax splice`. Each aligned read was assigned to a gene with FeatureCounts based on primary alignment. Reads with the same barcode and mapping to the same gene were UMI-collapsed with an error tolerance of 1 nt (L < 2). To determine cells above background, barcodes were ranked by UMI count. We applied the same percentile-based UMI count threshold as Cellranger (10X Genomics): 1/10th the UMI count of the barcode at the 99th percentile, plus an additional 10% of targeted cells. Reads were sorted by barcode and used to generate Seurat-compatible input files (barcodes.tsv, features.tsv, matrix.mtx) from each sample.

Cell filtering and clustering was performed using Seurat V3. Cells were filtered with the criteria: UMI count > 700, % mitochondrial genes < 20, and 0.4 < genes/UMIl < 0.8. Variable genes were identified using MeanVarPlot() with the following parameters: x.low.cutoff=0.0125, x.high.cutoff=3, y.cutoff=0.5. Principal component analysis was run with respect to these variable genes. Clustering was performed on the top 30 components using FindClusters() with a resolution of 0.6. Cell types were manually assigned using the top five positive and negative differentially-expressed genes per cluster.

### Gene and isoform quantification

Gene quantification in each cell was performed as previously described: barcoded reads were aligned to reference genome GRCm39 with Minimap2 and assigned with FeatureCounts to generate input matrices for downstream analysis.

To quantify transcripts, reads from each cell were aligned to the mouse reference transcriptome (GENCODE vM32, February 2023) using Minimap2 with the parameters: `--secondary=yes -N 100 -p 0.99 -ax map_ont.` The resulting alignment files were quantified using Salmon (v8) with the parameters: `salmon quant --ont -l U`. For each cell, this resulted in a Salmon output file columnating the total number of raw reads (`NumReads`) per isoform. Importantly, we took only the integer values from this column to make the data compatible with DESeq2, which operates on discrete count data. These count data were used to generate isoform count matrices for downstream analysis.

### Differential expression analysis

Differential expression analysis was performed using DESeq2’s likelihood ratio test (LRT) as recommended^53^. This analysis compares the fit of two linear models to each gene or isoform’s average expression across a variable of interest: a full model that includes the predictor variable of biological interest (i.e. cell type, experience, engram classification) and a reduced model that excludes this variable. To control for confounding effects of sex and batch on gene and isoform expression, we included both of these terms as fixed effects in all linear models.

For experience-variable differential expression analysis, the naive group (N) was arbitrarily assigned as baseline. Then for each gene or isoform feature, a separate coefficient was calculated for each pairwise comparison between baseline expression and average expression in the other experiences, resulting in a linear model equation with a coefficient (or slope) describing N vs C+ and another coefficient describing N vs U+. Because we included sex and batch as fixed effects, their respective coefficients were also calculated. The full model included all terms (experience, sex, and batch), while the reduced model only included sex and batch. To compare the fit of these models to the data, expression values were calculated for each cell using the full and reduced model equations. If the deviance across values predicted by the full model was smaller than for the reduced model (as determined by Analysis of Deviance (ANODEV)), the full model was considered an improved fit and the feature was considered experience-variable (adjusted p value < 0.05 after Benjamini-Hochberg correction (False Discovery Rate (FDR) ≤.05)). We used this differential expression framework for our cell-type and engram analyses, replacing the experience term in the full equation with cell type or engram status where appropriate.

To further classify experience-variable genes and isoforms by the experience(s) that upregulated them, we evaluated the log2 fold change for every pairwise comparison of average expression (C+ vs U+, C+ vs N, U+ vs N). A gene or isoform was classified as upregulated by an experience(s) if the log2 fold change of one or both comparisons was >.25.

## Data Availability

ONT scRNAseq files for each sample are available at NCBI Bioproject ID PRJNA1327322 (http://www.ncbi.nlm.nih.gov/bioproject/1327322).

## ACKNOWLEDGEMENTS

We would like to acknowledge Loyal Goff, Jean Fan, Greg Lewis, and C. Cavalier for their input, insight, intellectual and emotional support, as well as the mice who contributed their lives to this study.

This study was supported by funding from NIGMS (T32 GM007445 (Johns Hopkins BCMB)), NIH (R01 NS036715 (R.L.H.), R01 MH112152 (R.L.H.)), and NHGRI (R01 HG010538)(W.T.))

## Declaration of interests

W.T. has two patents (8,748,091 and 8,394,584) licensed to ONT and received reimbursement for travel, accommodation, and/or conference fees to speak at events organized by ONT.

## Notes

https://github.com/SheridanCavalier/2025_sc_iso_paper

