## Supplemental Material for "Single-cell long-read sequencing of the experience-induced transcriptome"

### SUPPLEMENTARY DATA

| Sample | Raw reads | % Demuxed | # Cells | Cell reads |
|---|---|---|---|---|
| **Oxford Nanopore Technologies (ONT) PromethION** | | | | |
| FC1 | 47,124,539 | 95 | 2153 | 6,250,785 |
| UC1 | 25,119,863 | 94 | 827 | 665,455 |
| NC1 | 43,889,067 | 95 | 1959 | 6,448,369 |
| FC2 | 22,813,281 | 91 | 1471 | 1,691,022 |
| UC2 | 17,606,346 | 93 | 1383 | 2,659,717 |
| NC2 | 15,431,986 | 94 | 1011 | 2,572,000 |
| FC3 | 44,500,123 | 90 | 1205 | 2,316,228 |
| UC3 | 47,589,095 | 92 | 1143 | 3,004,901 |
| NC3 | 50,915,289 | 91 | 1078 | 2,989,053 |
| FC4 | 27,119,682 | 93 | 1309 | 2,988,923 |
| UC4 | 36,286,978 | 93 | 738 | 1,803,771 |
| NC4 | 31,228,954 | 95 | 1285 | 4,252,938 |
| FC5 | 30,023,725 | 93 | 1110 | 3,017,142 |
| UC5 | 18,813,648 | 92 | 1110 | 1,766,903 |
| NC5 | 36,550,562 | 94 | 1299 | 4,019,512 |
| FC6 | 33,648,473 | 94 | 1098 | 3,458,023 |
| UC6 | 40,110,339 | 90 | 1887 | 4,431,455 |
| **Illumina MiSeq** | | | | |
| NC1 | 21,137,286 | 95 | 2445 | 13,612,412 |

**Table S1:** Raw reads from sequencer are demultiplexed against 10X whitelist. # Cells are determined by barcode rank. Cell reads column describes the final number of reads belonging to cell barcodes.

| Cluster | Cell type | Marker Gene | #Cells | Top Pos Genes in Cluster |
|---|---|---|---|---|
| **Oxford Nanopore Technologies (ONT) PromethION** | | | | |
| 0 | Dentate | *Prox1* | 608 | *Ncdn,Adcy1,Calb1,Grin2a,Nrgn,Cplx2* |
| 1 | Neuroblast | *Dcx* | 261 | *Igfbpl1,Sox11,Cd24a,Nnat,Gm3788,Gm9844* |
| 2 | Oligodendrocyte | *Mag* | 250 | *Ermn,Mag,Cldn11,Mog,Tspan2,Ugt8a* |
| 3 | Astrocyte | *Aldoc* | 225 | *Gja1,Sdc4,Aqp4,Ntsr2,Aldoc,Pla2g7* |
| 4 | Pyramidal | *Neurod6* | 147 | *Chrm3,Spink8,Wfs1,Sv2b,Pde1a,Ociad2* |
| 5 | Microglia | *Cx3cr1* | 137 | *Ctss,C1qb,C1qc,C1qa,Trem2,Fcrls* |
| 6 | Pericyte | *Vtn* | 77 | *Vtn,Higd1b,Atp13a5,Plac9a,Myl9,Kcnj8* |
| 7 | OPC | *Pdgfra* | 72 | *Pdgfra,Lhfpl3,Cacng4,C1ql1,Vcan,Cspg4* |
| 8 | Endothelial | *Cldn5* | 71 | *Cldn5,Slco1a4,Ly6c1,Flt1,Ly6a,Abcb1a* |
| 9 | Oligodendrocyte | *Mag* | 49 | *960013A20Rik,Bmp4,Rassf10,Gpr17,Enpp6,Kif19a* |
| 10 | Inhibitory | *Gad1* | 39 | *Slc32a1,Dlx6os1,Gad2,Dlx1,Dlx5,Pnoc* |
| 11 | Cajal Retzius | *Reln* | 23 | *Lhx1,Cacna2d2,Reln,Trp73,Lhx1os,Shisal2b* |
| **Illumina MiSeq** | | | | |
| 0 | Dentate | *Prox1* | 574 | *C1ql3,Pcp4,Ncdn,Olfm1,Calb1,Ncald* |
| 1 | Astrocyte | *Aldoc* | 266 | *Gja1,Aqp4,Ntsr2,PLa2g7,Clu,Aldoc* |
| 2 | Oligodendrocyte | *Mag* | 265 | *Mog,Mag,Cldn11,Ermn,Tspan2,Mal* |
| 3 | Neuroblast | *Dcx* | 218 | *Igfbpl1,Sox11,Cd24a,Neurod1,Nnat,Calb2* |
| 4 | Microglia | *Cx3cr1* | 189 | *C1qb,C1qa,C1qc,Ctss,Cx3cr1,Trem2* |
| 5 | Pyramidal | *Neurod6* | 179 | *Chrm3,4921539H07Rik,Pde1a,Spink8,Sv2b,Neurod6* |
| 6 | Dentate | *Prox1* | 164 | *Grin2a,Slit3,Kalrn,Dlgap2,Lrrtm4,Cacna1c* |
| 7 | Endothelial | *Cldn5* | 125 | *Cldn5,Flt1,Ly6c1,Ly6a,Slco1a4,Itm2a* |
| 8 | OPC | *Pdgfra* | 104 | *Pdgfra,Lhfpl3,Cacng4,Nxph1,C1ql1,Vcan* |
| 9 | Pericyte | *Vtn* | 80 | *Vtn,Atp13a5,Ndufa4l2,Kcnj8,Higdb,Myl9* |
| 10 | Neuroblast | *Dcx* | 76 | *Pclaf,Top2a,Eomes,Mki67,Ube2c,Lockd* |
| 11 | Oligodendrocyte | *Mag* | 56 | *9630013A20Rik,Bmp4,Enpp6,Rassf10,Gpr17,Bfsp2* |
| 12 | Inhibitory | *Gad1* | 53 | *Dlx6os1,Gad2,Slc32a1,Kcnmb2,Gm13629,Pnoc* |
| 13 | Pyramidal | *Neurod6* | 36 | *Stmn2,Malat1,Hapln4,Cck,Cnih2,Snrpn* |
| 14 | Cajal Retzius | *Reln* | 24 | *Cacna2d2,Reln,Trp73,Ebf3,Ndnf,Cd274* |
| 15 | Pericyte | *Vtn* | 18 | *Col1a1,Dcn,Slc6a13,Aox3,Lama1,Pcolce* |
| 16 | Dentate | *Prox1* | 18 | *Frmd3,Gm14372,Glra2,Fxyd7,Robo1,Bhle22* |

**Table S2:** Cell-type identification of clusters in Illumina and ONT single-cell data. Top six positive genes in each cluster by p value are shown, along with the expression marker used to assign cell type.

| | | % Assigned to Illumina Cell Type | | | | | | | | | | | |
|---|---|---|---|---|---|---|---|---|---|---|---|---|---|
| **ONT cell type** | **# Cells** | Astrocyte | CR | Dentate | Endothelial | Inhibitory | Microglia | Neuroblast | Olig | OPC | Pericyte | Pyramidal | ONT only |
| Astrocyte | 225 | 88.00 | 0.00 | 0.00 | 0.00 | 0.00 | 0.00 | 10.67 | 0.44 | 0.00 | 0.44 | 0.44 | 0 |
| Cajal Retzius (CR) | 23 | 0.00 | 100.00 | 0.00 | 0.00 | 0.00 | 0.00 | 0.00 | 0.00 | 0.00 | 0.00 | 0.00 | 0 |
| Dentate | 608 | 0.00 | 0.00 | 98.85 | 0.00 | 0.00 | 0.00 | 0.00 | 0.00 | 0.16 | 0.00 | 0.82 | 0.16 |
| Endothelial | 71 | 0.00 | 0.00 | 0.00 | 98.59 | 0.00 | 0.00 | 0.00 | 0.00 | 0.00 | 0.00 | 0.00 | 1.41 |
| Inhibitory | 39 | 0.00 | 0.00 | 0.00 | 0.00 | 92.31 | 0.00 | 0.00 | 0.00 | 0.00 | 0.00 | 2.56 | 5.13 |
| Microglia | 137 | 0.00 | 0.00 | 0.73 | 0.00 | 0.00 | 99.27 | 0.00 | 0.00 | 0.00 | 0.00 | 0.00 | 0.00 |
| Neuroblast | 261 | 0.00 | 0.00 | 7.66 | 0.00 | 0.38 | 0.00 | 90.80 | 0.00 | 0.38 | 0.00 | 0.38 | 0.40 |
| Olig | 250 | 0.00 | 0.00 | 0.00 | 0.00 | 0.00 | 0.00 | 0.00 | 98.40 | 0.80 | 0.00 | 0.40 | 0.40 |
| OPC | 121 | 0.00 | 0.00 | 0.00 | 0.00 | 0.00 | 0.00 | 0.00 | 0.00 | 100.00 | 0.00 | 0.00 | 0.00 |
| Pericyte | 77 | 0.00 | 0.00 | 0.00 | 2.60 | 0.00 | 0.00 | 0.00 | 0.00 | 0.00 | 97.40 | 0.00 | 0.00 |
| Pyramidal | 147 | 0.00 | 0.00 | 0.00 | 0.00 | 0.68 | 0.00 | 0.00 | 0.00 | 0.00 | 0.68 | 97.28 | 1.40 |

**Table S3:** Percentage of ONT barcodes by cell-type assigned to corresponding Illumina cell types.

| Sample | Experience | Time (min) | Age (days) | Sex | Batch |
|---|---|---|---|---|---|
| FC1 | Conditioned | 60 | 32 | F | A |
| UC1 | Unconditioned | 60 | 31 | F | A |
| NC1 | Naive | 0 | 31 | F | X |
| FC2 | Conditioned | 10 | 33 | F | B |
| UC2 | Unconditioned | 10 | 34 | M | B |
| NC2 | Naive | 0 | 30 | M | A |
| FC3 | Conditioned | 60 | 30 | M | C |
| UC3 | Unconditioned | 60 | 31 | M | C |
| NC3 | Naive | 0 | 32 | M | C |
| FC4 | Conditioned | 10 | 44 | M | D |
| UC4 | Unconditioned | 10 | 43 | F | D |
| NC4 | Naive | 0 | 33 | M | E |
| FC5 | Conditioned | 60 | 45 | F | D |
| UC5 | Unconditioned | 60 | 32 | M | E |
| NC5 | Naive | 0 | 31 | F | F |
| FC6 | Conditioned | 10 | 30 | M | F |
| UC6 | Unconditioned | 10 | 32 | F | F |

**Table S4:** Sample metadata.

| Cluster | Cell type | Marker Gene | #Cells | Top Pos Genes in Cluster |
|---|---|---|---|---|
| 3 | Astrocyte | *Aldoc* | 1472 | *Gja1,Aldoc,Clu,Slc1a3,Atp1a2,Plpp3* |
| 6 | CA1 | *Spink8* | 843 | *Neurod6,Ociad2,Rasgrp1,Wfs1,Spink8,Rprml* |
| 10 | CA2 | *Amigo2* | 581 | *Snhg11,Rian,Nrxn3,Meg3,Ryr3,Chrm3* |
| 16 | CA3 | *Bok* | 404 | *Selenow,Cck,Malat1,Cox8a,Stmn2,Scn1b* |
| 0 | Dentate_1 | *Prox1* | 2997 | *Calb1,Ncdn,Grin2a,Cplx2,Ptk2b,Nsf* |
| 4 | Dentate 2 | *Prox1* | 1320 | *Chgb,Scg2,Ntng1,Grin2a,Ncdn,Atp2b2* |
| 7 | Dentate_3 | *Prox1* | 858 | *Tpt1,Eif1,Ryr2,Gm10123,Kalrn,Meg3* |
| 8 | Endothelial | *Cldn5* | 688 | *Flt1,Ly6c1,Itm2a,Cldn5,Ly6a,Slco1a4* |
| 14 | Inhibitory | *Gad1* | 465 | *Gad2,Snhg11,Gad1,Nrxn3,Slc6a1,Dlx6os1* |
| 2 | Neuroblast | *Dcx* | 1624 | *Igfbpl1,Sox11,Gm9844,Nnat,Tmsb10,Tubb2b* |
| 1 | Olig_1 | *Mag* | 2155 | *Mog,Mag,Cldn11,Ermn,Trf,Tspan2* |
| 11 | Olig_2 | *Mag* | 526 | *Opalin,Mog,Mag,Ermn,Cldn11,Lpar1* |
| 9 | OPC | *Pdgfra* | 668 | *Pdgfra,Lhfpl3,Cspg5,Vcan,Ptprz1,Cacng4* |
| 5 | Microglia_1 | *Cx3cr1* | 1211 | *C1qa,C1qb,C1qc, Ctss, Cx3cr1,Fcer1g* |
| 13 | Pericyte_1 | *Vtn* | 516 | *Vtn,Rgs5,Plac9a,Flt1,Higd1b,Ly6c1* |
| 12, 15 | Progenitor | *Sox11* | 924 | *Gm10282,Mdk,Hmgb2,Hmgb3,Lmnb1* |
| 18 | Pericyte_2 | *Vtn* | 121 | *Igfbp2,Dcn,Slc7a11,Col1a2,Itih5,Vtn* |
| 20 | Microglia_2 | *Cx3cr1* | 56 | *Pf4,Mrc1,Lyz2,Dab2,Ctsc,Ms4a7* |
| 22 | Olig_3 | *Mag* | 42 | *Cldn11,Mog,Ermn,Ptgds,Opalin,Mag* |
| 23 | Olig_4 | *Mag* | 24 | *Spata17,Aqp4,Slc13a1,Col24a1,Gja1,Gm22299* |
| 17 | CR | *Reln* | 40 | *Ndnf,Reln,Cacna2d2,Nhlh2,Shisa12b,Slc17a6* |
| 19 | Unassigned 1 | | 151 | *Myh11,Acta2,Tpm2,Myl9,Crip1,Plac9a* |
| 21 | Unassigned 2 | | 116 | *Tmem212,Ccdc153,Meig1,Rsph1,Dynlrb2,Aebp1* |

**Table S5:** Cell-type identification of clusters in the combined ONT single-cell dataset. Top six positive genes in each cluster by p value are shown, along with the expression marker used to assign cell type.

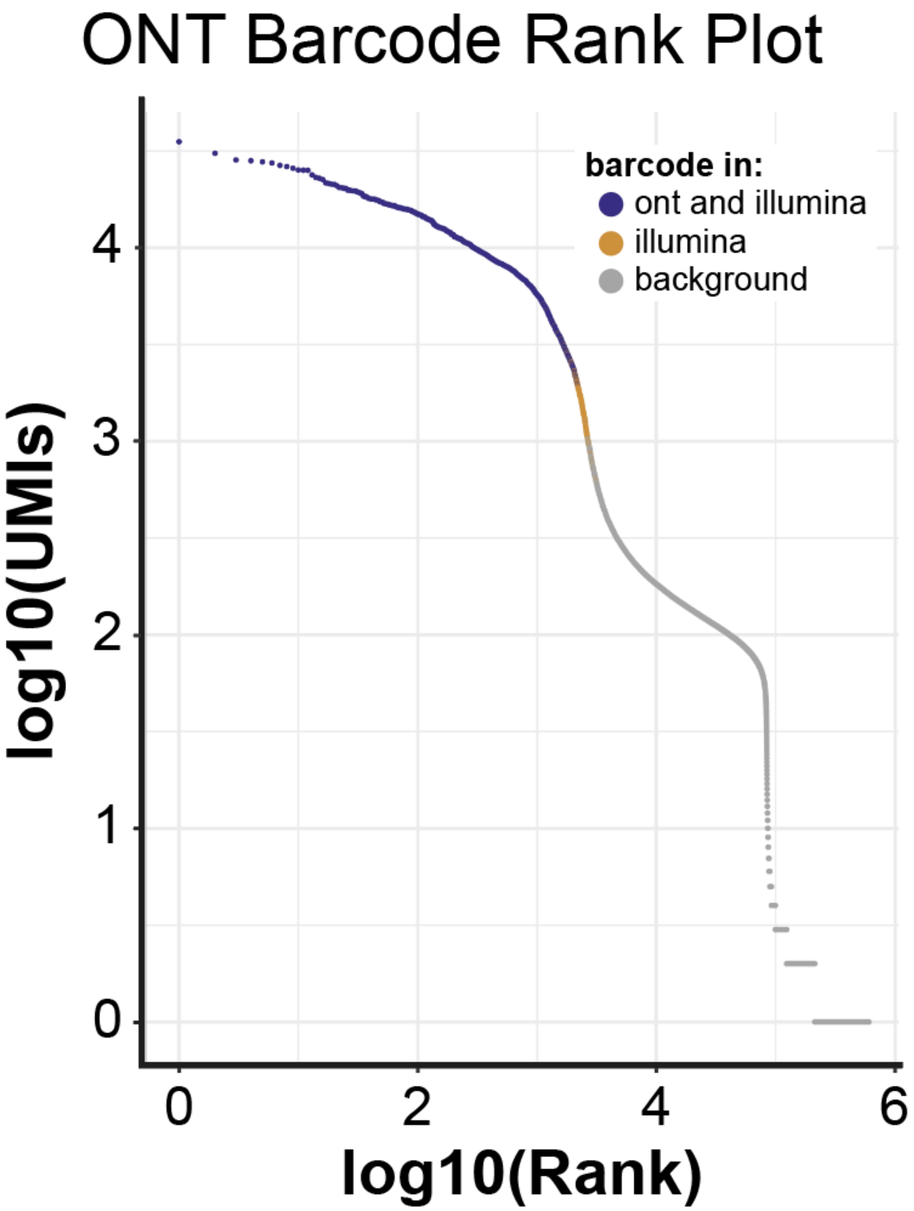


**Fig S1:** ONT barcode rank plot. All barcodes recovered in the ONT dataset were ordered by UMI count. 'ONT and Illumina' barcodes (blue) meet the UMI count threshold required to qualify as cells in both datasets, while 'Illumina' barcodes (orange) only qualify as cells in the Illumina data.

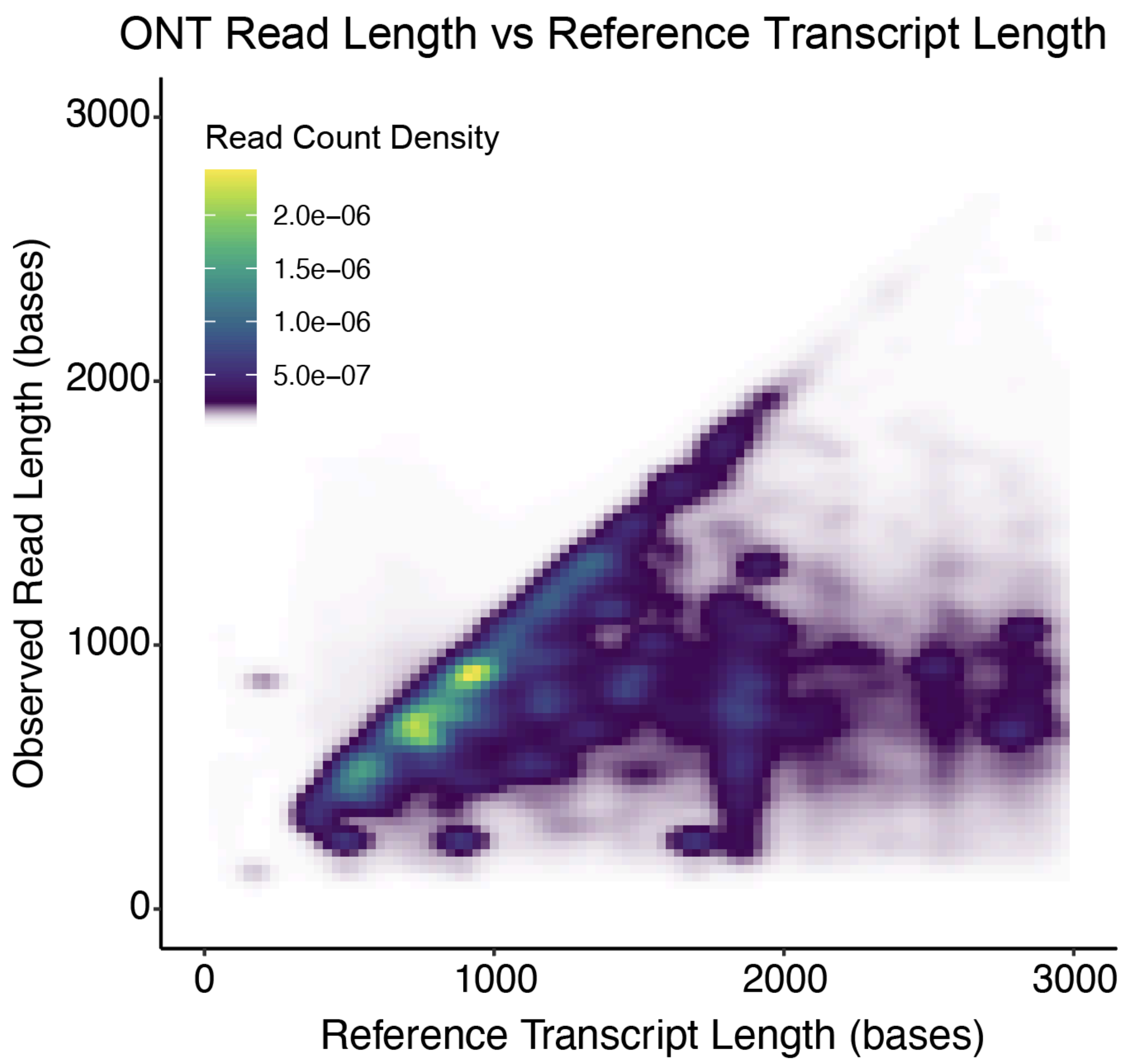


**Fig S2:** Observed insert lengths from cell-derived ONT reads compared to reference transcript lengths (assigned by primary transcriptome alignment). Lengths of reference transcripts come from GENCODE VM32.

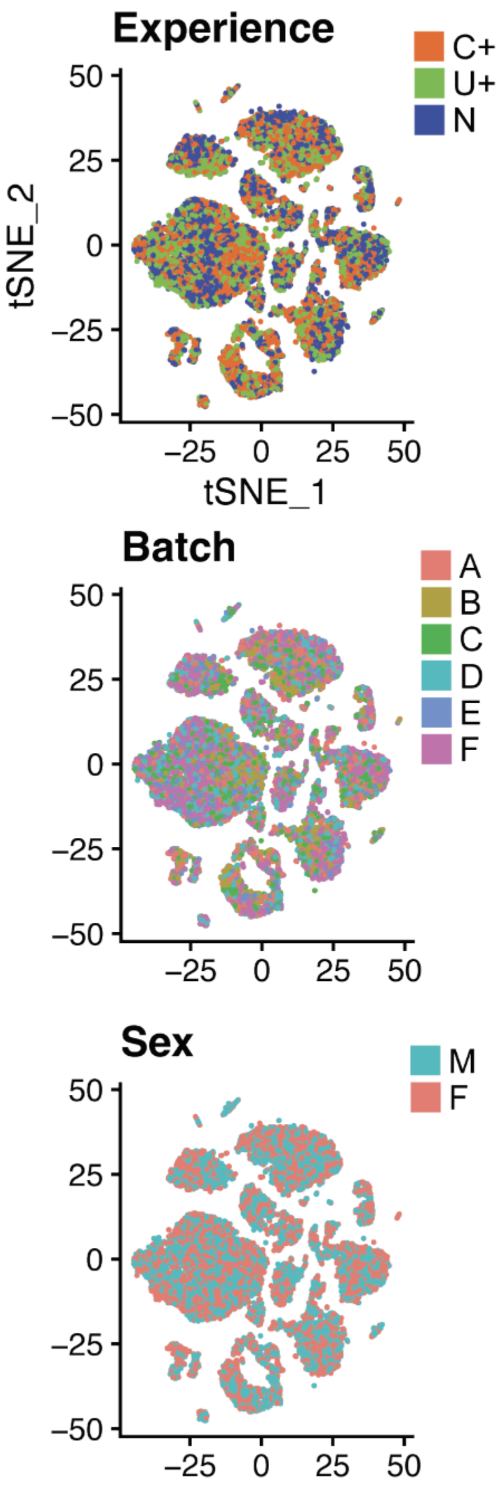


**Fig S3:** tSNEs from combined ONT hippocampal samples (n=16) colored by experience (top), batch (center), and sex (bottom).

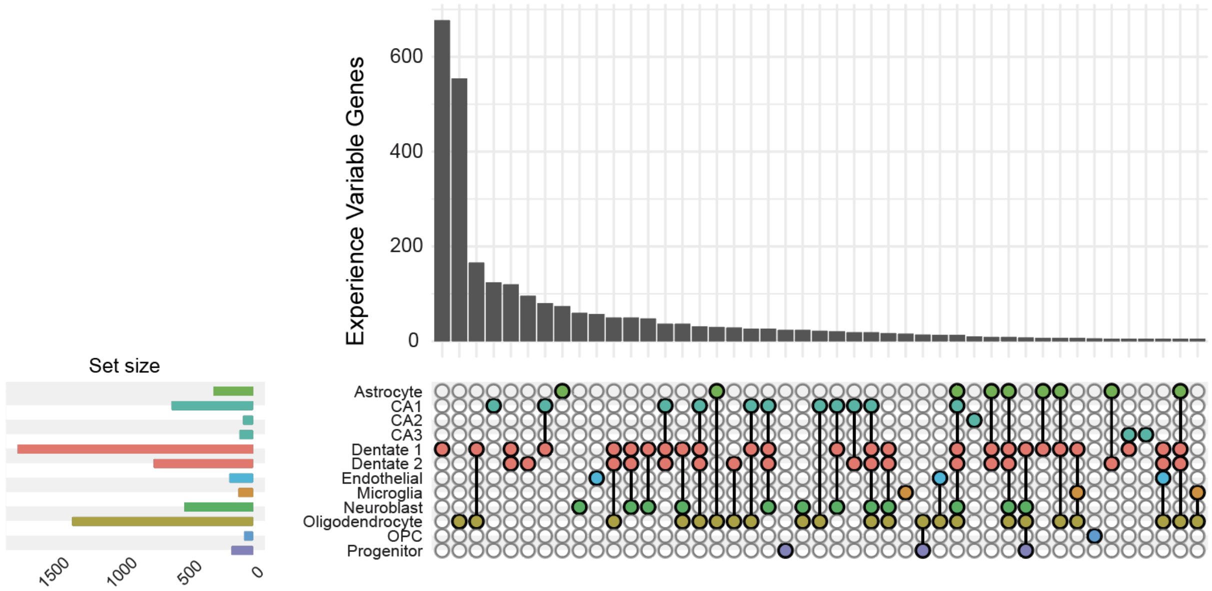


**Fig S4:** Upset plot of experience-variable genes across hippocampal cell types. Intersections are filtered to contain >4 genes.

GO Biological Process Terms (2023)

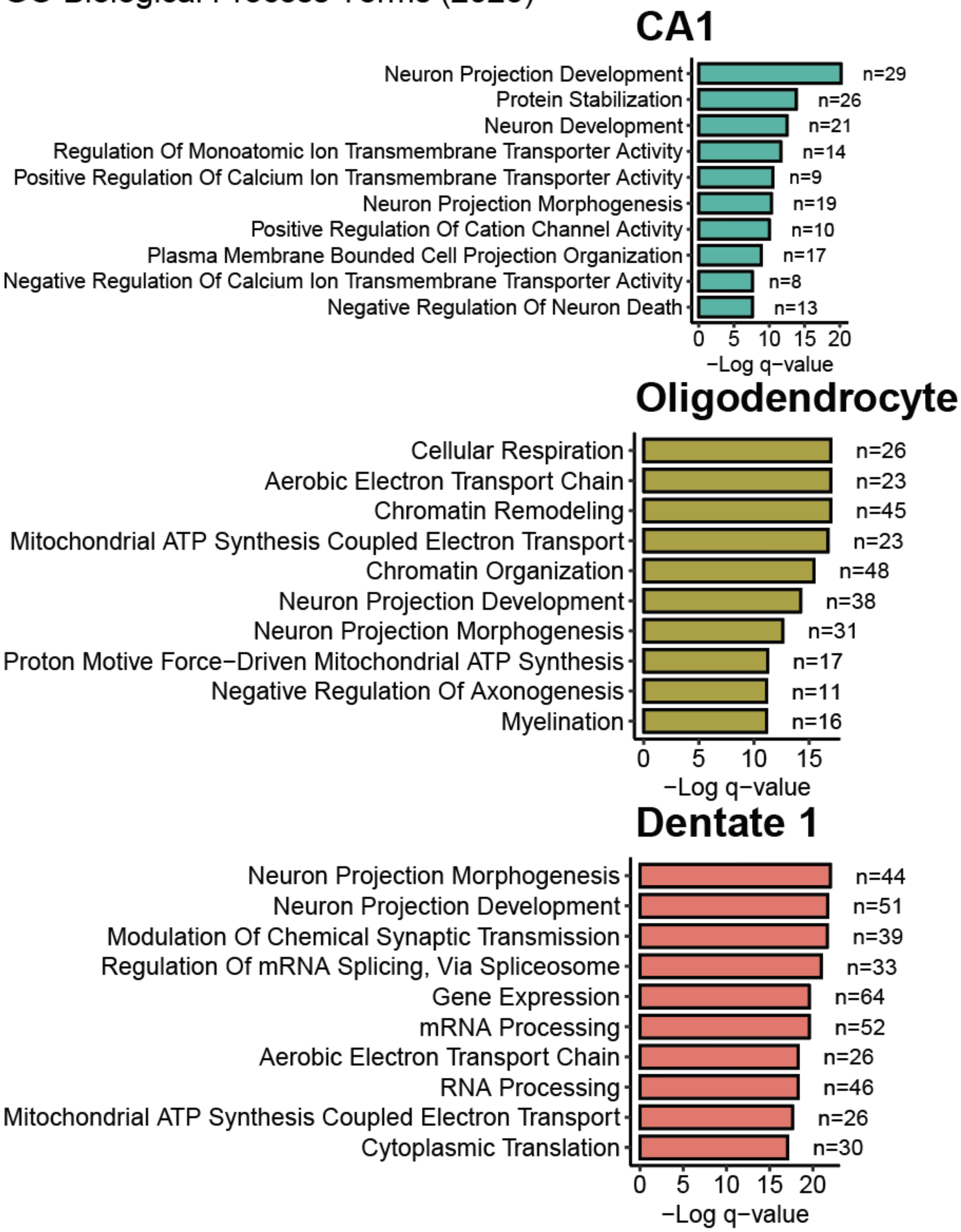


**Fig S5:** GO enrichment analysis results for experience-variable genes in CA1 neurons, Dentate 1 neurons, or Oligodendrocytes. GO enrichment analysis was performed against the GO Biological Process (2023) database. The top 10 terms (by q value) are shown for each cell type.

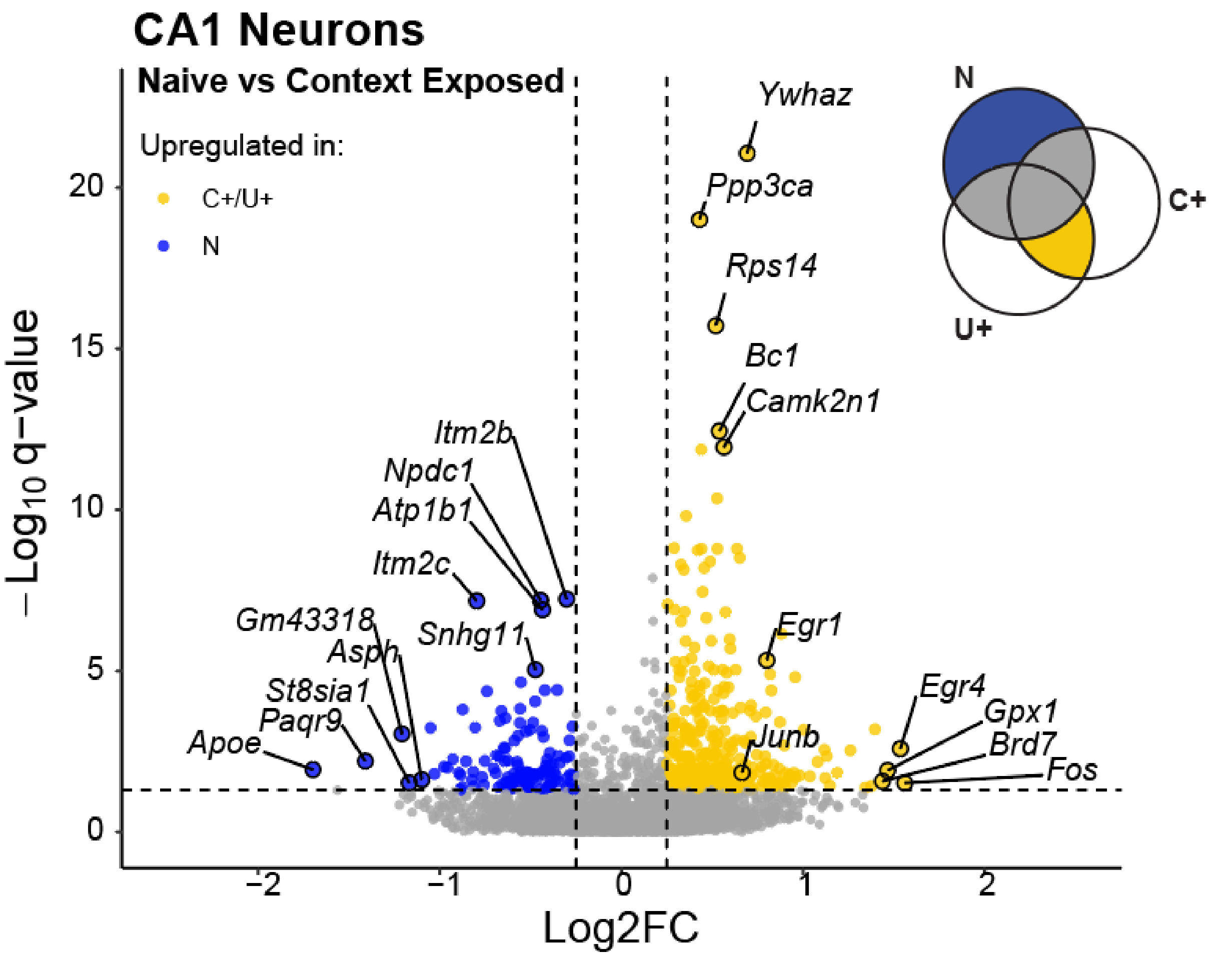


**Fig S6:** Volcano plot of differentially-expressed genes between context-exposed and naive CA1 neurons. For each gene, expression in C+ and U+ cells was averaged and compared with expression in N cells resulting in the log2 fold change values shown. Q values are from DESeq2 analysis. The top five genes by significance and log2 fold change per condition are labeled in addition to *Egr1* and *Junb*.

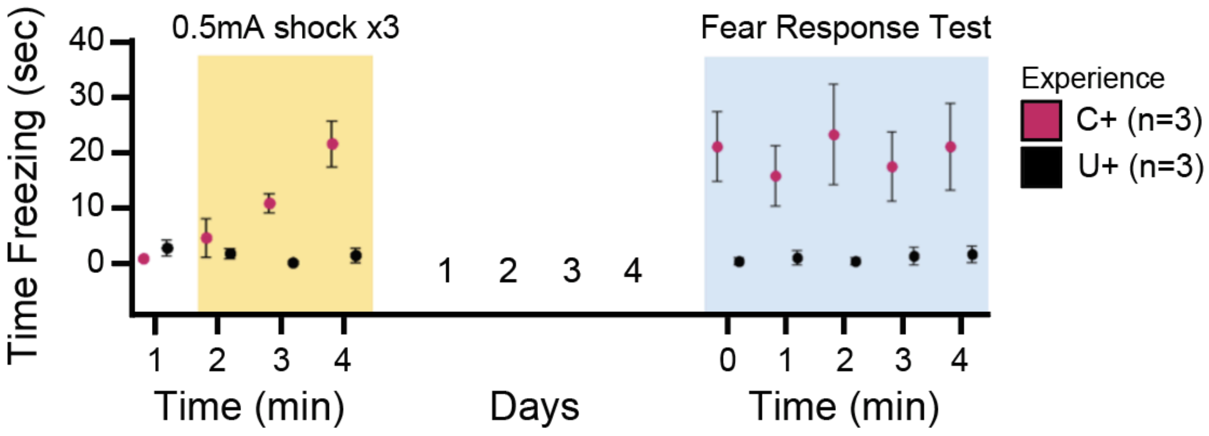


**Fig S7:** Conditioned fear response test from fear conditioned (C+) vs unconditioned (U+) littermates. During context exposure, C+ mice received three 1sec, 0.5 mA footshocks while U+ mice received no footshocks. C+ and U+ mice were then reintroduced to the context in absence of footshocks four days later. Time spent freezing (sec) is quantified during context exposure and again during context re-introduction. C+ mice spend significantly more time freezing during context reintroduction than U+ littermates, indicating contextual learning.

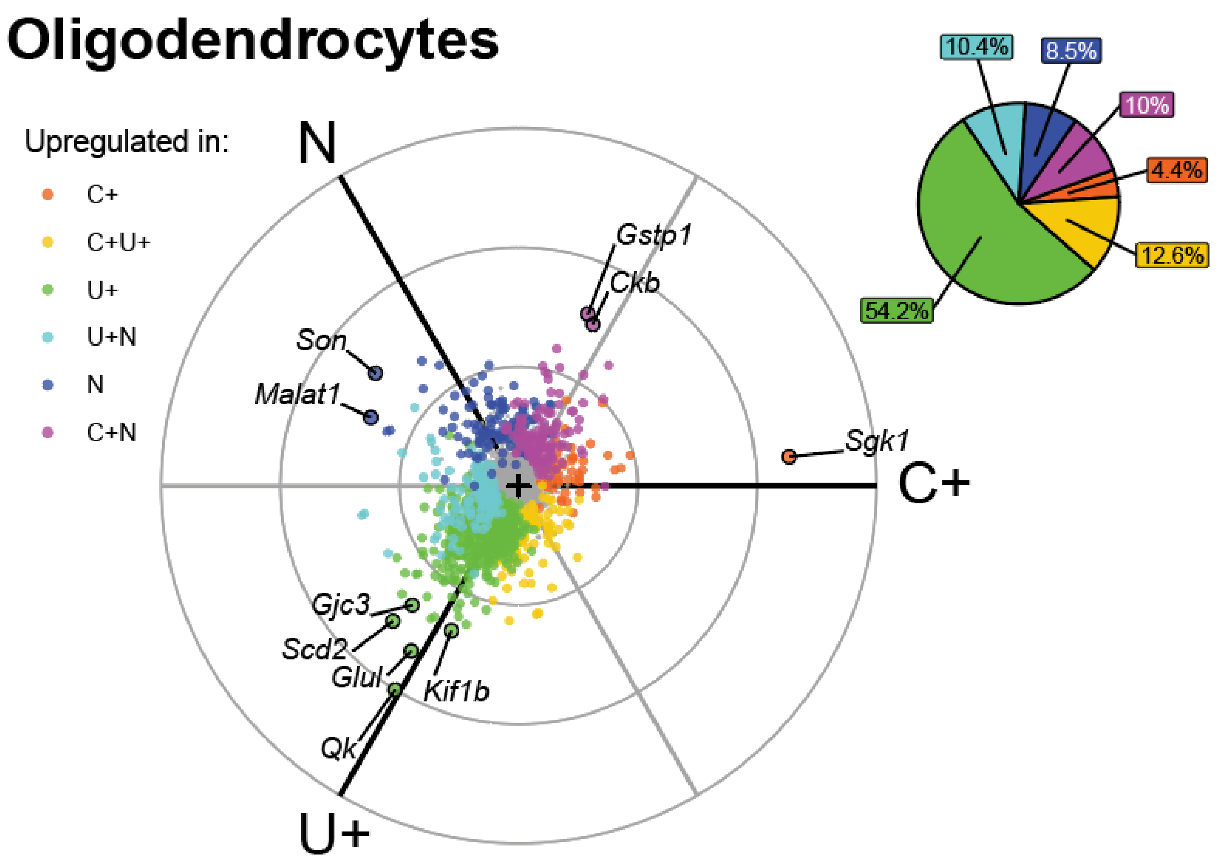


**Fig S8:** Radial expression plot of genes in oligodendrocytes. Significantly experience-variable genes (q value < .05, FDR ≤ .05) are colored by the experience(s) in which they were determined to be upregulated (log2FC > .25). Top ten genes by *r* value are labeled. (inset) Percentage of total genes upregulated by each experience.

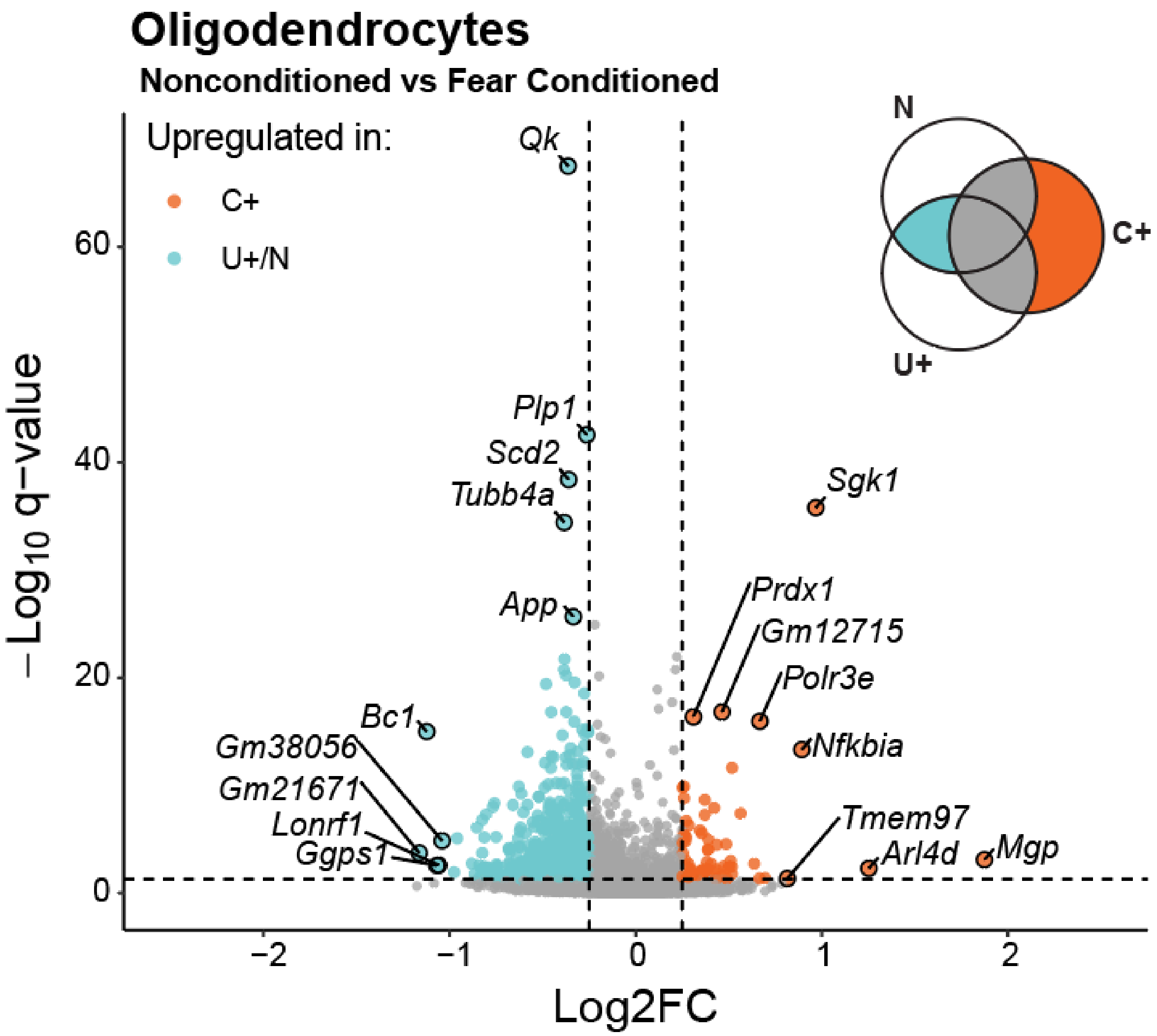


**Fig S9:** Volcano plot of differentially-expressed genes between fear conditioned and nonconditioned oligodendrocytes. For each gene, expression in C+ cells was compared with the averaged expression in U+ and N cells, resulting in the log2 fold change values shown. Q values are from DESeq2 analysis.The top five genes per condition by significance and log2 fold change are labeled.

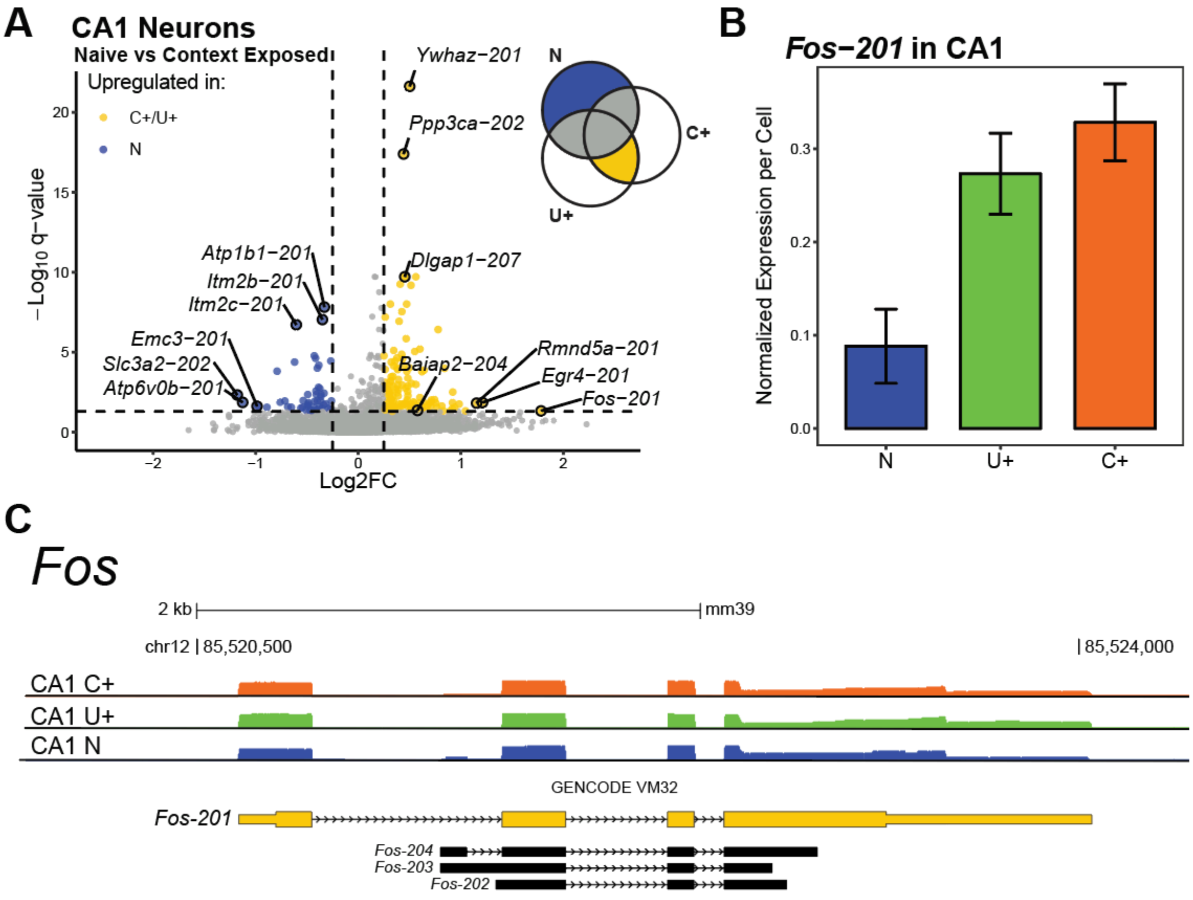


**Fig S10: A.** Volcano plot of differentially-expressed isoforms between context-exposed and naive CA1 neurons. For each isoform, expression in C+ and U+ cells was averaged and compared with expression in N cells resulting in the log2 fold change values shown. Q values are from DESeq2 analysis.The top three isoforms per condition by significance and log2 fold change are labeled in addition to *Baiap2-204*. **B.** Average normalized expression of isoform *Fos-201* per cell, within each experience. Error bars represent standard error. **C.** Coverage histogram tracks from CA1 reads aligning to *Fos*, aggregated by experience and visualized using UCSC Genome Browser. Tracks are individually scaled to maximum coverage. Highlighted (yellow) is the exposure-upregulated isoform *Fos-201*.

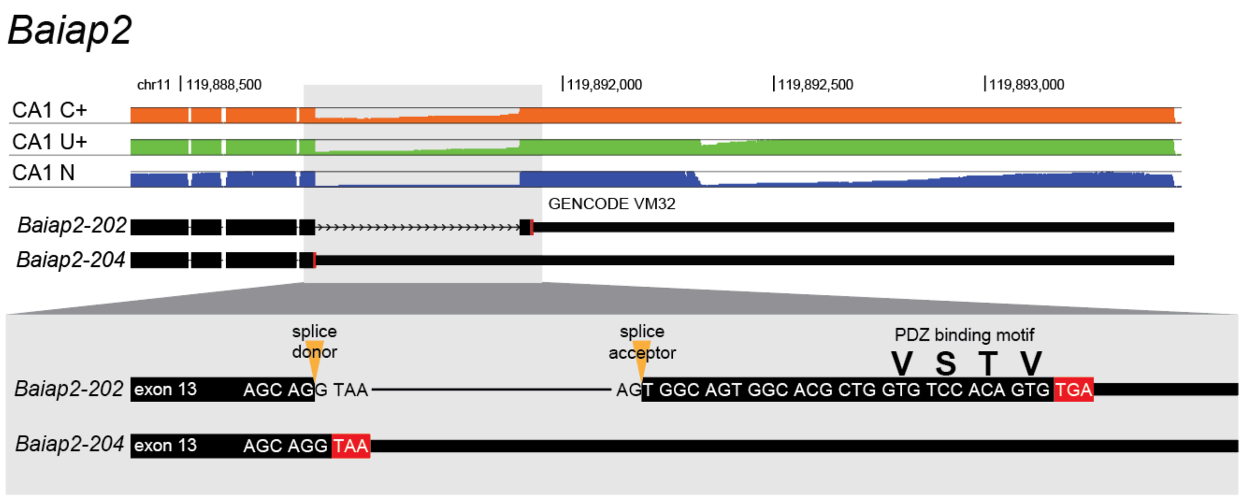


**Fig S11:** Coverage histogram tracks from CA1 reads aligning to *Baiap2*, aggregated by experience and visualized using UCSC Genome Browser. Tracks are configured to show only exonic features and are scaled to 50 reads. Highlighted is the alternative splicing decision that yields the two isoforms, with *Baiap2-202* encoding the PDZ binding motif (VSTV) and exposure-upregulated *Baiap2-204* excluding it.

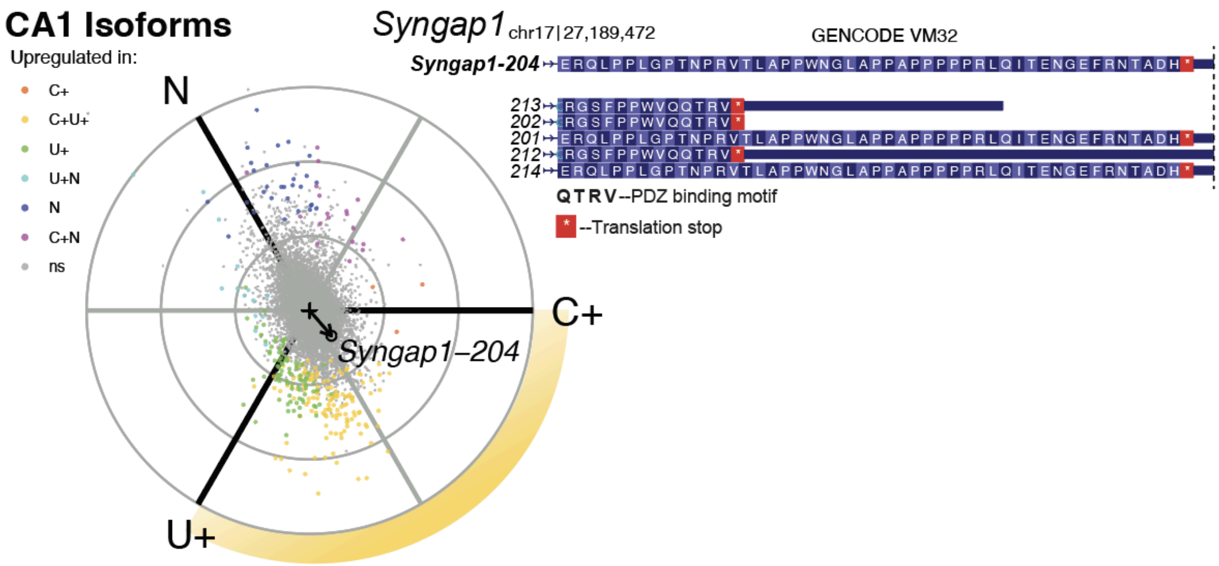


**Fig S12:** (left) Radial expression plot of transcript isoforms in the CA1, with *Syngap1-204* labeled. (right) Amino acid sequences for the final exons of *Syngap1* isoforms (GENCODE VM32), visualized in UCSC Genome Browser. *Syngap1-204* yields an isoform of the protein that does not include the C-terminal PDZ binding motif (QTRV).
